# Temperature compensation in a small rhythmic circuit

**DOI:** 10.1101/716761

**Authors:** Leandro M. Alonso, Eve Marder

## Abstract

Temperature affects the conductances and kinetics of the ionic channels that underlie neuronal activity. Each membrane conductance has a different characteristic temperature sensitivity, which raises the question of how neurons and neuronal circuits can operate robustly over wide temperature ranges. To address this, we employed computational models of the pyloric network of crabs and lobsters. We employed a landscape optimization scheme introduced previously (Alonso and Marder, 2019) to produce multiple different models that exhibit triphasic pyloric rhythms over a range of temperatures. We use the currentscapes introduced in (Alonso and Marder, 2019) to explore the dynamics of model currents and how they change with temperature. We found that temperature changes the relative contributions of the currents to neuronal activity so that rhythmic activity smoothly slides through changes in mechanisms. Moreover, the responses of the models to extreme perturbations—such as gradually decreasing a current type—are often qualitatively different at different temperatures.

## I. Introduction

Dynamic biological systems depend on many interacting nonlinear processes that together produce complex outputs. In the nervous system, neuronal activity requires the coordinated activation and inactivation of many inward and outward currents, mediated by a variety of different ion channels. Temperature influences all biological processes, to a greater or lesser degree. This poses an inherent difficulty for neuronal signaling: if the currents involved in neuronal and network dynamics are differentially temperature-dependent a system that is well-tuned to work at one temperature may not function at a different temperature (Caplan et al., 2014; O’Leary and Marder, 2016; Tang et al., 2010). Nonetheless, many ectothermic animals have neurons and circuits that do function well over an extended temperature range (Robertson and Money, 2012). It then becomes important to understand how and to what extent this can occur.

The effects of temperature on channel function and neuronal activity have been studied extensively for many years (Taylor and Kerkut, 1958). Increasing temperature generally results in increases of channel maximal conductances and faster activation/inactivation rates (Frankenhaeuser and Moore, 1963). But, importantly ionic channels of different types are affected by temperature to different extents: each of these processes has a different Qio (Schauf, 1973; Kukita, 1982; Ruff, 1999; Tang et al., 2010). The effect of temperature on neuronal intrinsic excitability such as voltage and current thresholds can be diverse (Sjodin and Mullins, 1958; Guttman et al., 1962, 1966; Fitzhugh, 1966). For example, temperature experiments (18 – 35°C) in identified neurons in locust showed reversible changes in spike amplitude and duration of spikes (Heitler et al., 1977), and studies in the jump neural circuit of grasshopers showed that neural excitability is differently affected by temperature across neuronal types (Abrams and Pearson, 1982).

There are also examples of neuronal and circuit processes that are relatively temperature-compensated. For example, the *f/I* curves in both in molluscan and locust neurons are not substantially affected by small temperature changes (6 – 8°C) (Connor, 1975; Heitler et al., 1977). A recent study showed that the *f/I* curves of auditory sensory neurons in grasshoppers remains largely unaffected by temperature changes over a range (21 – 29°*C*) (Roemschied et al., 2014). Temperature compensation also takes place the behavioral level. For example, a recent study in hunting archerfish showed that the duration of the two major phases of the C-start—a fast escape reflex which follows prey release—were temperature compensated (Krupczynski and Schuster, 2013). The question then arises of how these behaviors are preserved when the currents in the cells and the cells’ intrinsic excitability properties are differentially modified by temperature.

Here we explore these issues using computational models of the pyloric network—a subnetwork within the stomatogastric ganglion (STG) of crustaceans (Marder and Bucher, 2007; Maynard, 1972). The pyloric rhythm is a triphasic motor pattern that consists of bursts of action potentials in a specific sequence. This behavior is robust and stable, and the cells and their connections are well-characterized. Temperature compensation in the pyloric network has been explored experimentally by Tang et al. (2010, 2012) and Soofi et al. (2014). These studies show that as temperature increases the frequency of the pyloric rhythm increases but the phases of the cycle at which each cell is active remain approximately constant. Additional work on the pacemaker kernel of the pyloric network (three cells connected by gap junctions that burst synchronously) showed that temperature increases the frequency of these bursts (*Q*_10_ ≈ 2), but their duty cycle (the burst duration in units of the period) stays approximately constant (Rinberg et al., 2013).

Temperature compensation was explored in computational models of the pacemaking kernel (Soto-Trevino et al., 2005) by Caplan et al. (2014) and O’Leary and Marder (2016). They showed that it was possible to find multiple sets of *Q*_10_ values for different membrane conductances processes, so that the duty cycle of the cells remained constant as their bursting frequency increased. In this work we build on the results in Caplan et al. (2014) and implemented temperature sensitivity in a model of the pyloric network (Prinz et al., 2004). We show that in these models there are multiple sets of maximal conductances and temperature sensitivities that give rise to temperature compensated rhythms, and that they can reproduce much of the phenomenology previously reported (Tang et al., 2010, 2012; Soofi et al., 2014; Haddad and Marder, 2018). In addition, we explored how the dynamics of the currents are modified to sustain the correct activity at each temperature. We performed this study for 36 different models and found that in all cases, the contributions of the currents to the activity can be significantly different across temperatures. The currents are not simply scaled up but instead become reorganized in a complex way: for instance, a current that is important for burst termination at 10°*C* may no longer play that role at 25°*C*. The contribution of a given current to a neuronal process can be partially replaced by another. As a consequence of this, the stability properties of the activity can be different at different temperatures. Therefore, a second perturbation for which the system is not compensated, may trigger qualitatively different responses in individual circuits depending on the temperature. These results provide a plausible hypothesis for why interactions between temperature and a second perturbation are often observed experimentally (Haddad and Marder, 2018; Ratliff et al., 2018).

## II. Results

### A. Duty cycle and phase compensation in model pyloric networks

The pyloric rhythm is produced by the periodic sequential activation of three neuronal types: the pyloric dilator (PD) neurons which are electrically coupled to the the anterior burster (AB) neuron forming a pace-making kernel, the lateral pyloric (LP) neuron and five to eight pyloric (PY) neurons. We modified the model of the pyloric circuit in Prinz et al. (2004) to include temperature sensitivity. Figure 1A shows a schematic representation of the model pyloric network studied here (diagram adapted from Prinz et al. (2004)). The model network consists of three cells, each modeled by a single compartment with eight voltage-gated currents as in previous studies (Golowasch and Marder, 1992; Buchholtz et al., 1992; Goldman et al., 2001). The synaptic connections are given by seven chemical synapses of two types as in Prinz et al. (2004). In this model the pace-making kernel is aggregated into a single compartment *AB* – *PD* (here we refer to this compartment as PD). Following Liu et al. (1998), each neuron has a sodium current, *I_Na_*; transient and slow calcium currents, *I_CaT_* and *I_CaS_*; a transient potassium current, *I_A_*; a calcium-dependent potassium current, *I_KCa_*; a delayed rectifier potassium current, *I_Kd_*; a hyperpolarization-activated inward current, *I_H_*; and a leak current *I_leak_*. The traces in Figure 1A show a solution of this model for one set of maximal conductances *G*. The traces exhibit a triphasic pyloric rhythm that consists of the sequential bursting of the *PD, LP* and *PY* cells. This pattern remains approximately periodic so we can measure the phases of the cycle at which each cell is active, as indicated by the color labels.

**FIG. 1.**
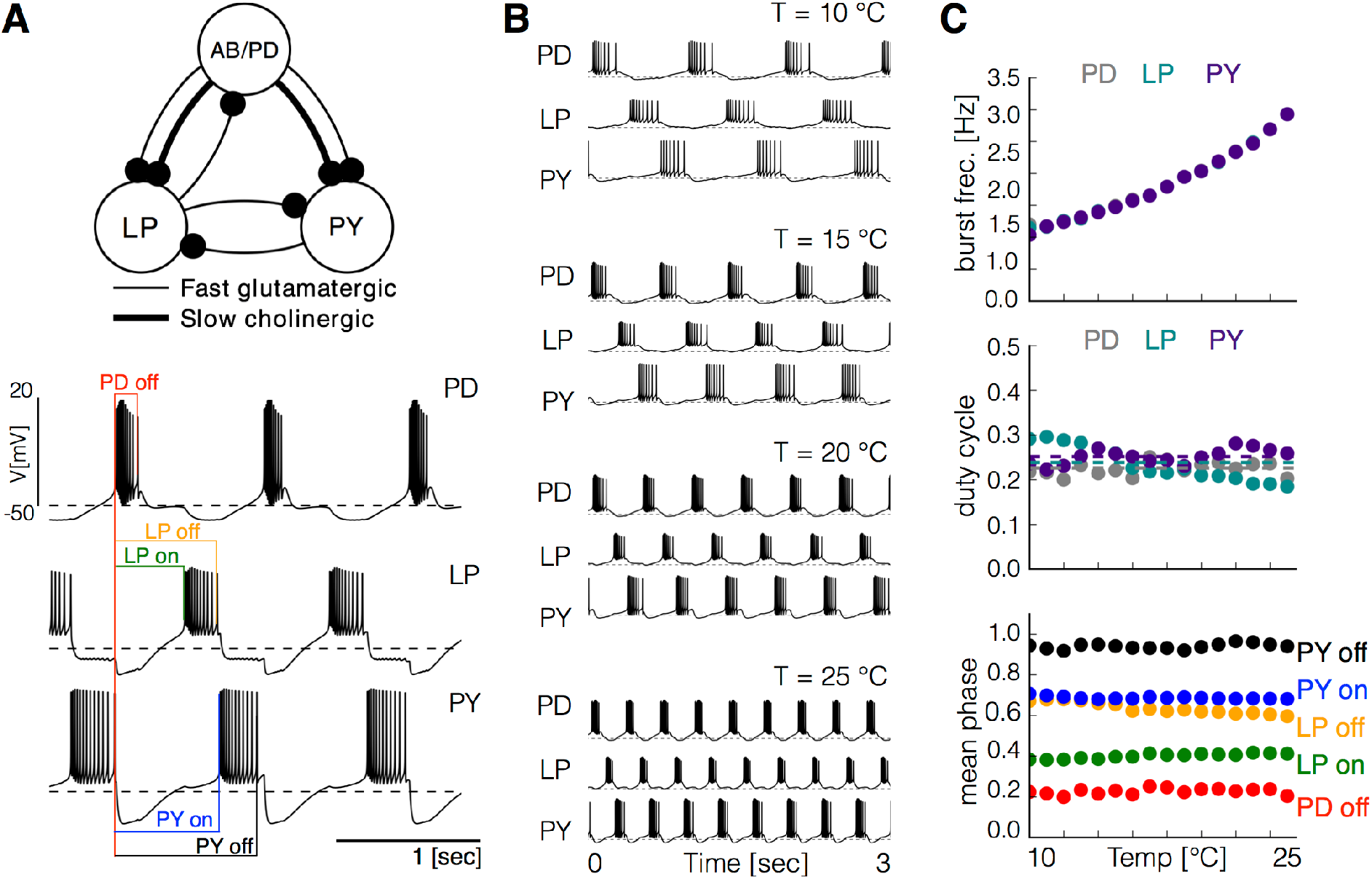
Temperature compensation in a model pyloric network. **A** Schematic diagram of the model pyloric network in Prinz et al. (2004). The three cells interact via seven chemical synapses of two types. The traces below show a representative solution that exhibits a triphasic rhythm: the activity is approximately periodic and the cells burst in a specific sequence: *PD-LP-PY*. **B** Activity of a temperature compensated model network at different temperatures 10°*C* – 25°*C*. As temperature increases the frequency of the rhythm increases but the duty cycle of the cells—the burst duration in units of the period—remains approximately constant. **C** The panels show average values over 30 seconds for 16 values of temperature between 10°*C* – 25°*C*. Top: average burst frequency of each cell. Middle: average duty cycle of each cell (cell type indicated in colors). Bottom: average phases of the cycle at which bursts begin and terminate. As temperature increases these phases remain approximately constant.

Temperature sensitivity was modeled for each conductance as in Caplan et al. (2014) and O’Leary and Marder (2016) (see Methods). We increased each conductance by an Arrhenius type factor 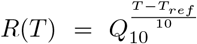 (with *Q*_10_ ≥ 1 and *T_ref_* = 10°*C*) and also increased the rates of the channel kinetics *τ* in similar fashion. We assume that *Q*_10_*G*__ ∈ [1,2] for the maximal conductances and *Q*_10_*τ*__ ∈ [1,4] for the timescales as these ranges are consistent with experimental measurements of these quantities. Because the properties of each channel type are expected to be similar across cells we assume the *Q*_10_ values for any given current are the same across cell and synapse types. Different values of the maximal conductances *G* can produce an innumerable variety of activities. The space of solutions of the model—the different patterns produced for each set of maximal conductances *G*—is formidably complex. This is in part due to the nonlinear nature of neuronal dynamics but also because the number of conductances that need to be specified is large (*dim*(*G*) =8 × 3 + 7 = 31). The picture is more complicated as we consider the effect of temperature because it affects both the conductances and the time scales of the different processes (14 in total). Therefore in the models, changes in temperature correspond to a coordinated change—a path—in a 45-dimensional parameter space. The complexity of this problem led us to confine our study to specific questions suggested by biological observations over a permissible temperature range.

In the crab, as temperature is increased pyloric activity remains triphasic—the cells fire in the same order and at the same relative phases—while the pyloric network frequency increases by a 2 to 3 fold factor over a 15°*C* range (Tang et al., 2010). For this to happen in the model, there should be many regions in parameter space for which the activity is triphasic at different frequencies and paths in parameter space—sequences of values of *G* and *τ*—that interpolate these regions. One of the main results in this work is that such paths do exist and that temperature compensation is possible in this model. Finding these temperature compensated networks is nontrivial: it requires the specification of 8 × 3 + 7 = 31 maximal conductances *G* and 24 *Q*_10_ values. Because it is virtually impossible to find these sets of parameters at random we employed a landscape optimization scheme described previously (Alonso and Marder, 2019) and adapted it to this particular scenario (see Methods). An alternative approach was described in O’Leary and Marder (2016). We first searched for sets of maximal conductances *G* that displayed triphasic rhythms at control temperature (10°*C*). Inspired by physiological recordings of the pyloric rhythm of crabs and lobsters we targeted control activities with a network frequency of 1*Hz*, duty cycle ≈ 20% for *PD*, and duty cycle ≈ 25% for *LP* and *PY* (Bucher et al., 2006; Hamood et al., 2015). We then selected 36 of these models and for each of them we searched for *Q*_10_ values so that the activity is preserved over a temperature range. We found that, regardless of the maximal conductances *G*, it was always possible to find multiple sets of *Q*_10_ values that produced a temperature compensated pyloric rhythm.

Figure 1B shows the activity of one example network at several temperatures. As temperature increases the frequency of the rhythm increases but the phases of the cycle at which each cell is active remains approximately constant. This is consistent with the experimental results reported by Tang et al. (2010). Figure 1C shows the same analysis as in Tang et al. (2010) performed on this particular model. We simulate the model at several temperatures over the working range and compute the average duty cycle, frequency and phases of the bursts. While the burst frequency increases with *Q*_10_ ≈ 2, the duty cycle of the cells, and the phases of the onsets/offsets of activity, remain approximately constant. Figure 1SA supplemental shows the same analysis performed on three different models. In all cases the duty cycle and the phases stay approximately constant while the frequency increases. Figure 1SB shows the values of the maximal conductances *G* and *Q*_10_ for each of these models.

### B. Effect of temperature on membrane potential

Experimental studies on other systems reported effects of temperature on resting membrane potentials (Klee et al., 1974), spiking thresholds, and amplitudes of spikes (Heitler et al., 1977). In the models studied here, some temporal properties of the activity remain approximately constant over a temperature range but the precise shape of the waveforms can change. To inspect how the membrane potential changes over the working temperature range (10°*C* – 25°*C*) we computed the distribution of *V* of each cell for *R* = 101 values of temperature. Temperature changes features of the voltage waveforms that result in changes in these distributions. Figure 2A shows example traces and the result of this analysis for the *LP* cell in one model. The membrane potential changes with temperature in several ways: the spike amplitude decreases and the spike threshold depolarizes. At each temperature, we computed the distribution of *V* and used a gray scale to indicate the fraction of time that the cell spends at each voltage value (y-axis). The color lines match features of the voltage waveforms at each temperature, with features in the distributions on the right. The distributions permit visualizing changes in the waveform as the control parameter is changed (Alonso and Marder, 2019).

**FIG. 2.**
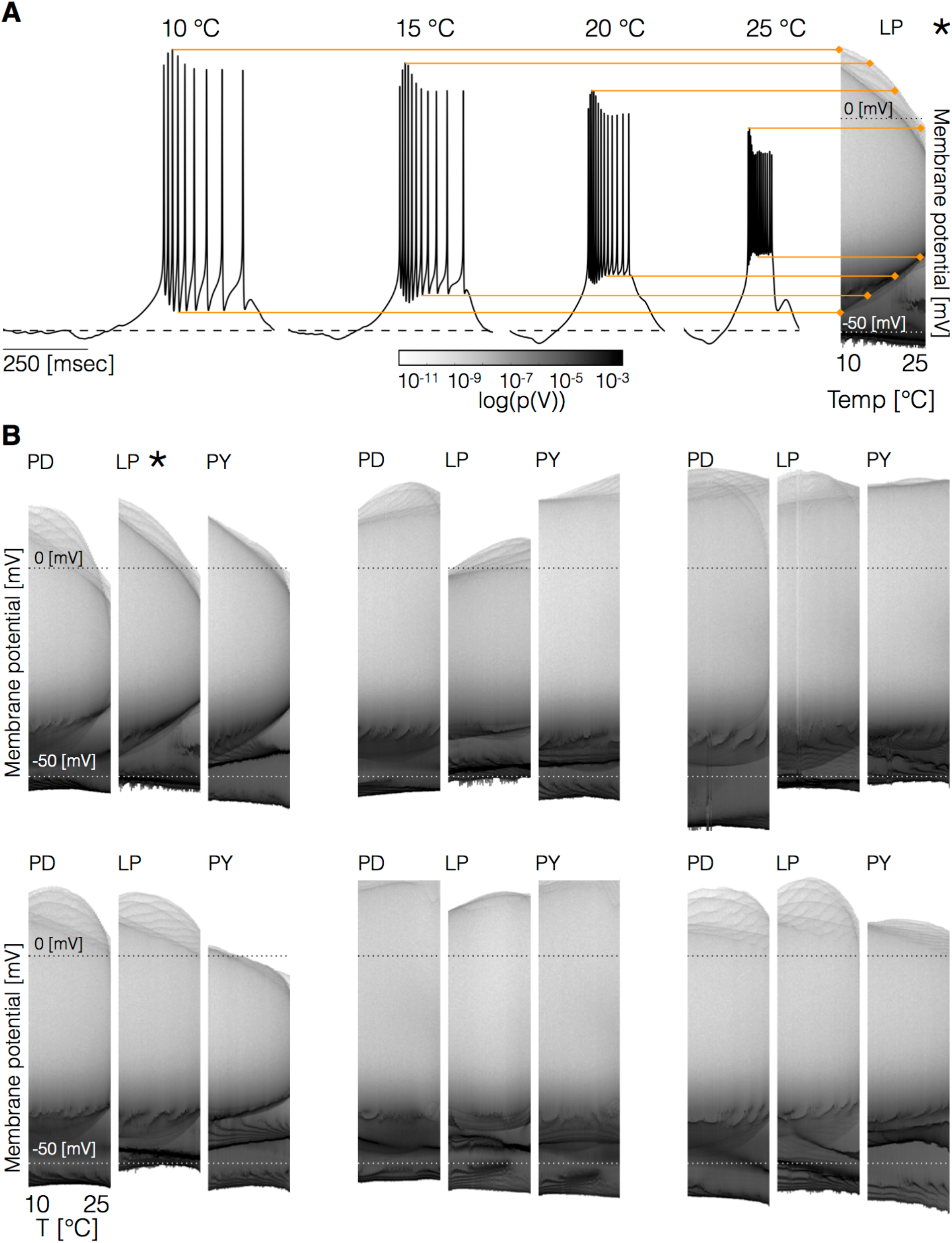
Changes in membrane potential over temperature. All cells remain bursting with similar duty cycles over the working temperature range but the waveforms of the membrane potential exhibit changes. **A** (left) Representative traces of the *LP* cell in one model at different temperatures. (right) Membrane potential distribution at each temperature. The gray scale indicates the proportion of time the cell spends at that potential. The distributions facilitate inspecting how features such as the total amplitude of the oscillations and spiking thresholds change with temperature. **B** The panels show the membrane potential distributions of each cell over temperature for 6 models. The oscillation amplitudes, spiking thresholds and other features of the sub-threshold activity show clear differences across models.

Figure 2B shows the membrane potential distributions of each cell for 6 models. In all cases the waveforms show visible differences across temperatures. In some cases the peak voltages of the spikes—indicated by the upper envelope of the distributions—remains relatively constant while in other cases it changes. There is considerable variability in the amplitudes of the oscillations across models. The distributions also show how some features of the sub-threshold activity change with temperature. In particular, in the models on the top, we observe an increase in the spiking thresholds while in the models in the bottom the spiking threshold remains relatively constant or decreases. In addition, note that cells can either become more depolarized or more hyperpolarized as temperature increases.

### C. Spiking patterns during temperature ramps

In the models, as temperature is increased the values of the maximal conductances increase and the time scales of the channels kinetics decrease (the processes become faster). While we showed previously that the duty cycle remains—on average—approximately constant, the spiking patterns of the cells do show discernible differences across temperatures. To show this we subjected the models to linear temperature ramps from 10°*C* to 25°*C* over 60 minutes, as depicted in Figure 3A, and recorded the spike times. Figure 3B shows the inter-spike interval (*ISI*) distributions for each cell over the working temperature range (10–25°*C*). Notice that at each temperature, there is one large *ISI* that corresponds to the inter-burst interval and many small *ISI* values, which correspond to the spike intervals within a burst. As temperature is increased the spiking pattern changes in complicated ways and the largest *ISI* decreases monotonically, consistent with the overall increase in bursting frequency of the cells.

**FIG. 3.**
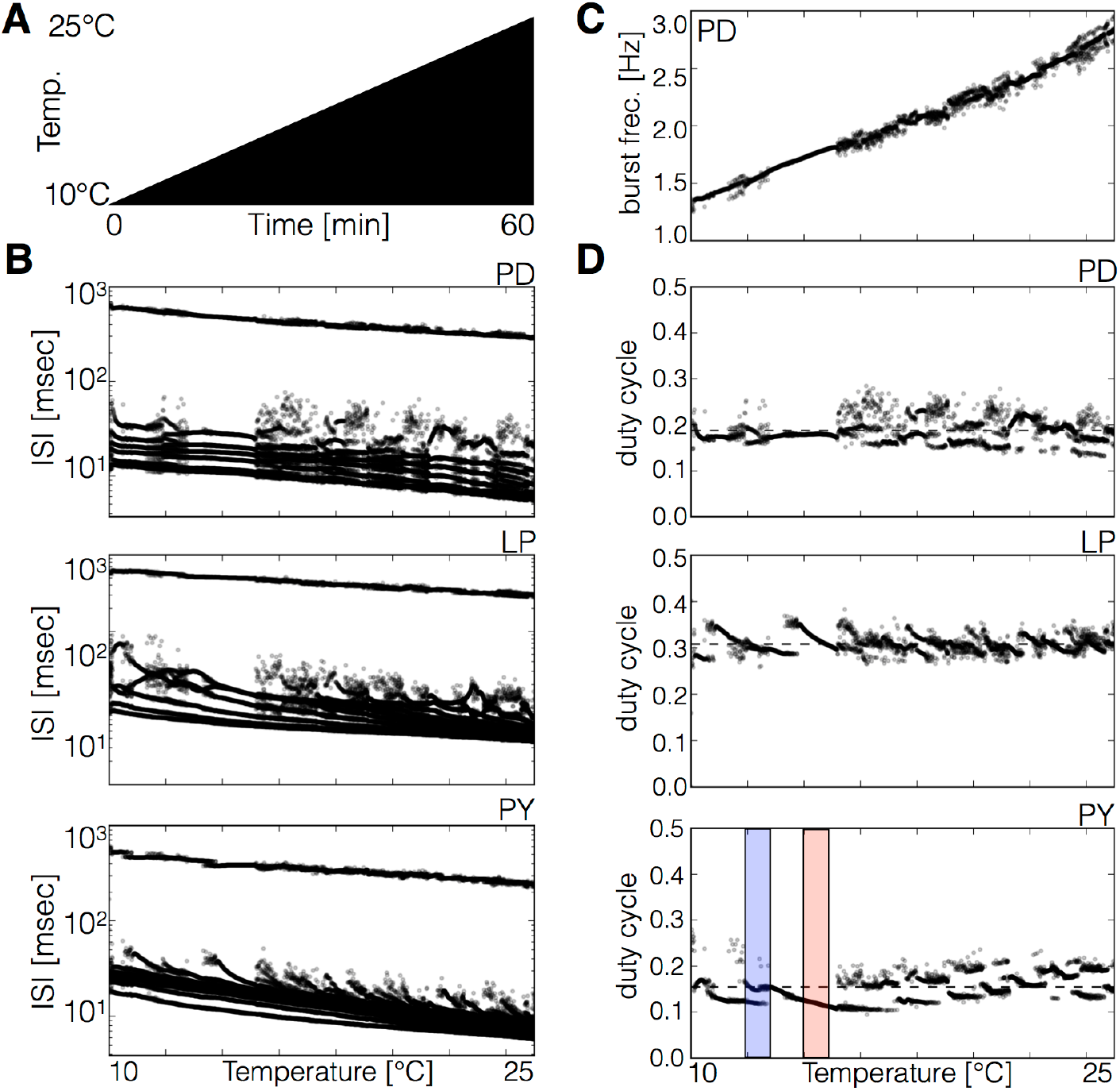
Spiking patterns during temperature ramps. The spiking patterns of the cells exhibit interesting dependences with temperature. **A** Schematic representation of the temperature ramp used in these simulations. Temperature was increased linearly between 10°*C* and 25°*C* over 60 minutes. **B** ISI of each cell over temperature (y-axis is logarithmic). The spiking patterns consist of one large ISI corresponding to the inter-burst interval (≥ 200*msec*) and several smaller values produced by spikes within bursts. There are ranges of temperature over which the ISI takes almost the same values and the patterns result in nearly identical sets of points, and there are ranges where there is more variability. **C** Instantaneous frequency of each burst of the *PD* cell. **D** Duty cycle of each burst in each cell. While the duty cycle remains approximately constant on average, there are ranges of temperature for which the duty cycle alternates between two or more values (pink and blue shaded boxes).

We define a spike to be a part of a burst if it occurs within 200msec of another spike. Using this criterion we can compute the duty cycle and instantaneous frequency of each burst. Figure 3C shows the instantaneous burst frequency of every burst in the *PD* cell. As temperature increases the frequency of the bursts increases and there is little variability from burst to burst. Figure 3D shows the distribution of duty cycles for each cell over the working temperature range. Notice that different values of temperature result in slightly different spiking patterns which in turn result in different values of the duty cycle. For some temperatures, the duty cycle is nearly identical in every burst resulting in a single point in the y-axis. For example, this is the case for the temperatures indicated by the pink box in the PY cell. There are temperatures for which the duty cycle takes two values (blue box in PY cell) and there are temperatures for which the duty cycle differs noticeably from burst to burst, and produce a filled region in the diagrams. In most if not all cases, this difference corresponds to a last spike that may be absent in some bursts. While this is unsurprising it is interesting that it happens in some temperature ranges and not in others.

Although in all models the average duty cycle stays approximately constant as temperature is changed, the duty cycle distributions can show marked differences across models: different maximal conductances and different *Q*_10_ values produce different patterns. We focus on these spiking patterns because they can be compared directly to both intracellular and extracellular experimental recordings. Figure 4 shows the duty cycle distributions of the *LP* cell over the working range for all 36 models. We chose to show this cell because the biological network contains 2 *PD* cells and 5 *PY* cells but only one *LP* cell which makes comparisson to experiments simpler. In all cases there are temperatures for which the duty cycle takes almost exactly one value, ranges where it takes two values, and ranges where multiple values are produced. Overall Figure 4 shows that these patterns can be diverse across models and provides a notion of how variable the bursting patterns of the cells can be across temperatures and individuals.

**FIG. 4.**
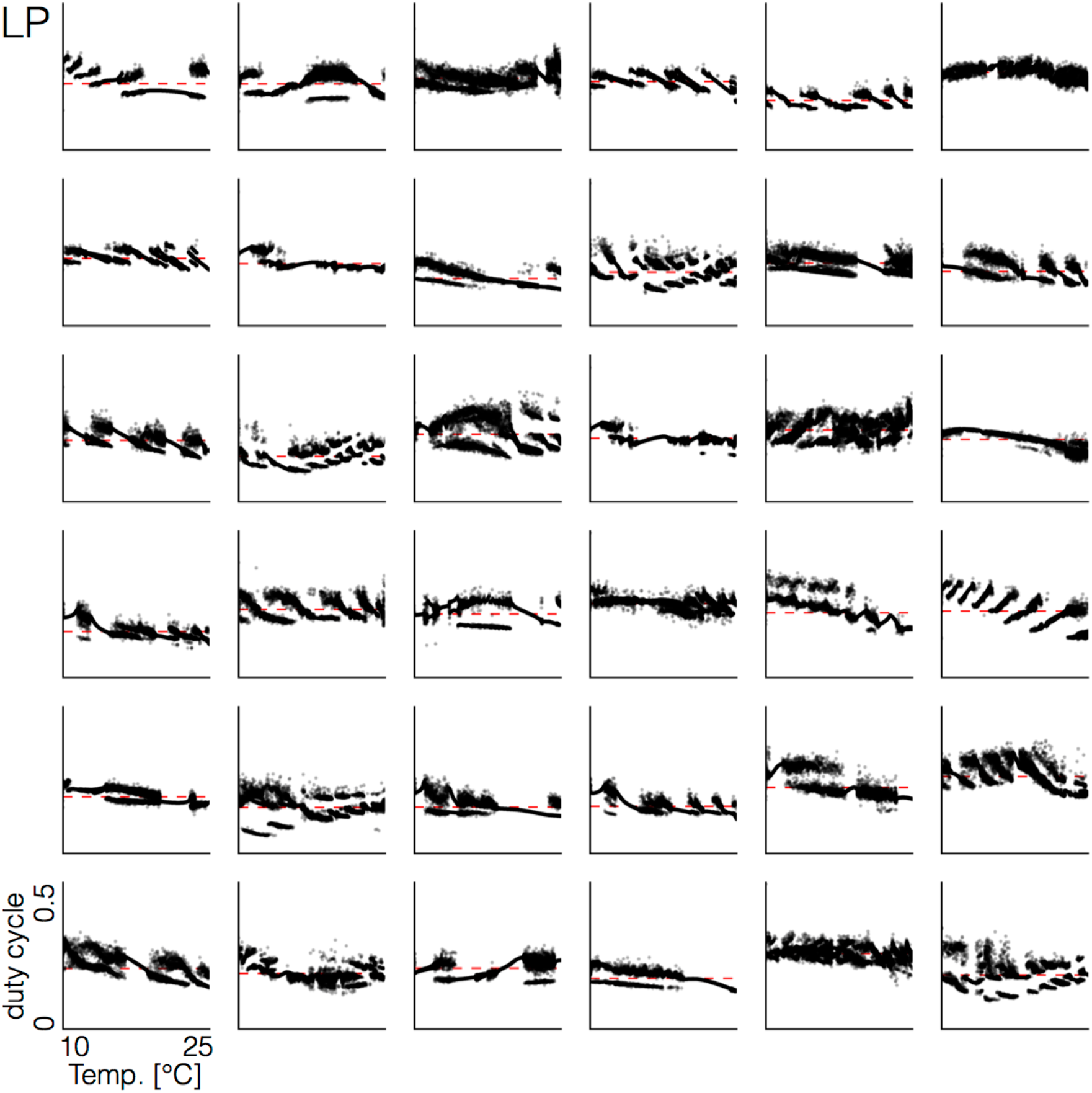
Duty cycle distributions of all models. There is significant variability across models in the way the duty cycle distributions change with temperature. The figure shows the distributions of duty cycle of the *LP* cell for all 36 models during the same temperature ramp as in Fig. 3. The precise shape of these patterns depends on the values of the maximal conductances and temperature sensitivities.

### D. Hysteresis and multistability

Hysteresis and multistability are often observed in conductance based models of neuronal activity (Malashchenko et al., 2011; Marin et al., 2013), and the network studied here also exhibits such features. We explored hysteretic behavior in the models by increasing temperature linearly from 10°*C* to 35°*C* in 30 minutes and then decreasing it back to 10°*C* at the same pace. Figure 5 shows the spiking patterns for two models during one such temperature ramp (schematized on top). The ISI distributions show normal bursting activity up to a critical temperature where the model becomes quiescent. The quiescent state remains stable as temperature continues to increase and eventually recovers activity during the decreasing ramp, at a lower temperature than when it stopped spiking during the increasing ramp. The temperature at which the model switches from spiking to quiescence is different in each side of the ramp. This implies that there is a range of temperatures where the model can be either quiescent or spiking, and thus is multistable.

**FIG. 5.**
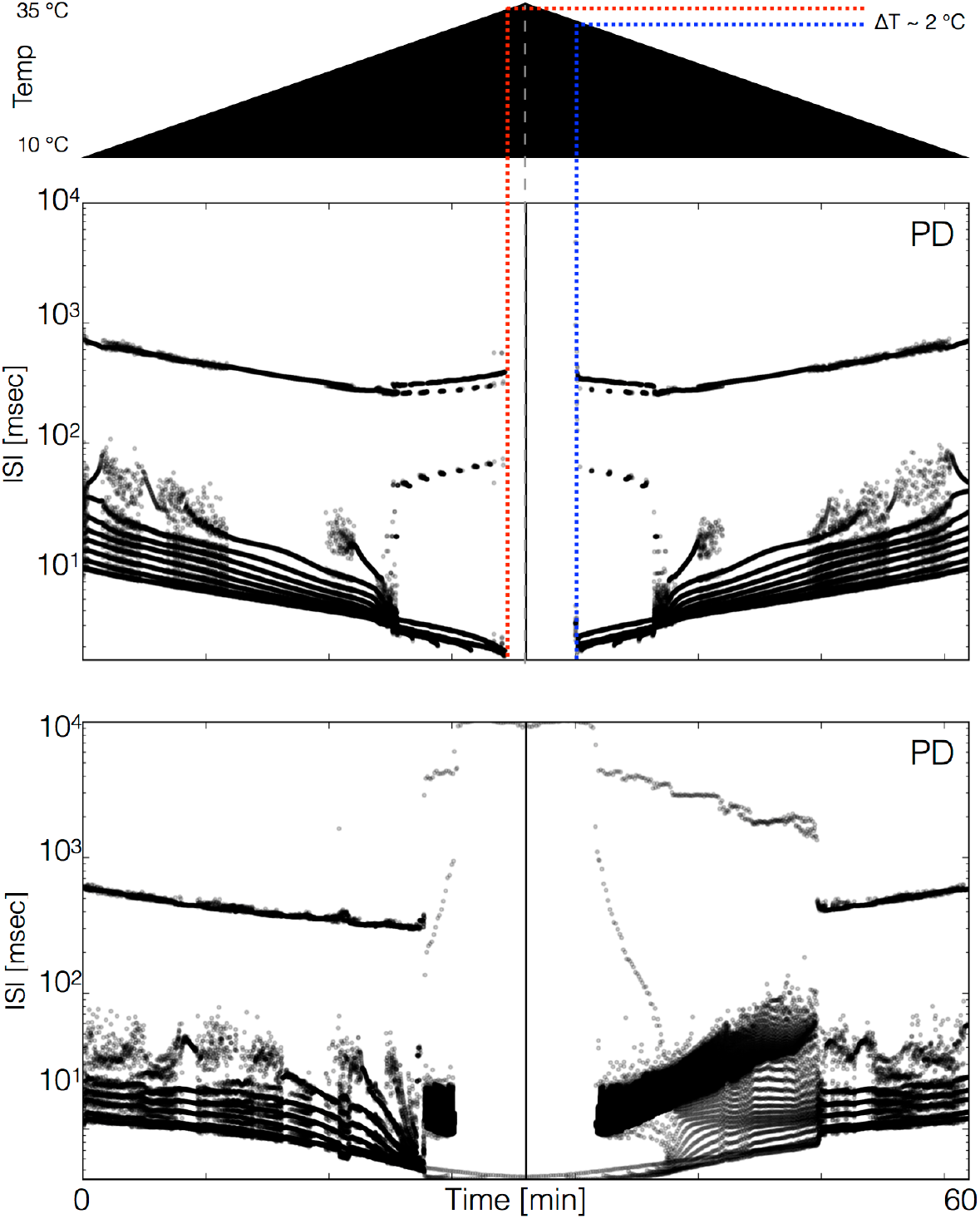
Hysteresis and multistability. We ramped the temperature in the models from 10°*C* to 35°*C* and then back to 10°*C* in a symmetric fashion. The figure shows a schematic representation of the temperature ramp on the top. The bottom part shows the ISI distributions of the *PD* cell over time for two different models. **(top)** The cell ceases to produce spikes before 600 secs, which is also before reaching 35°*C*, and remains quiescent until temperature is ramped back down. The temperature at which the cell resumes spiking is about ≈ 2°*C* lower than the temperature at which it became quiescent. This indicates that the system is multi-stable over that temperature range. **(bottom)** Hysteresis is more evident in this model. The spiking patterns during the down ramp differ visibly from those during the up ramp over a wide temperature range.

Multistability is ubiquitous in these models. The bottom panel in Fig. 6 shows a different model in which these features are more salient. At high temperatures (for times between 25 and 35 minutes) this model produces *ISI* values as long as ≈ 1*sec* and it remains producing spikes at all temperatures. The spiking patterns differ noticeably on the up and down ramps. These observations suggest that at many temperatures, there are multiple stable attractors that correspond to the triphasic rhythm but differ slightly in their spiking patterns. In many cases these differences come down to the precise timing of the last spike of a burst.

**FIG. 6.**
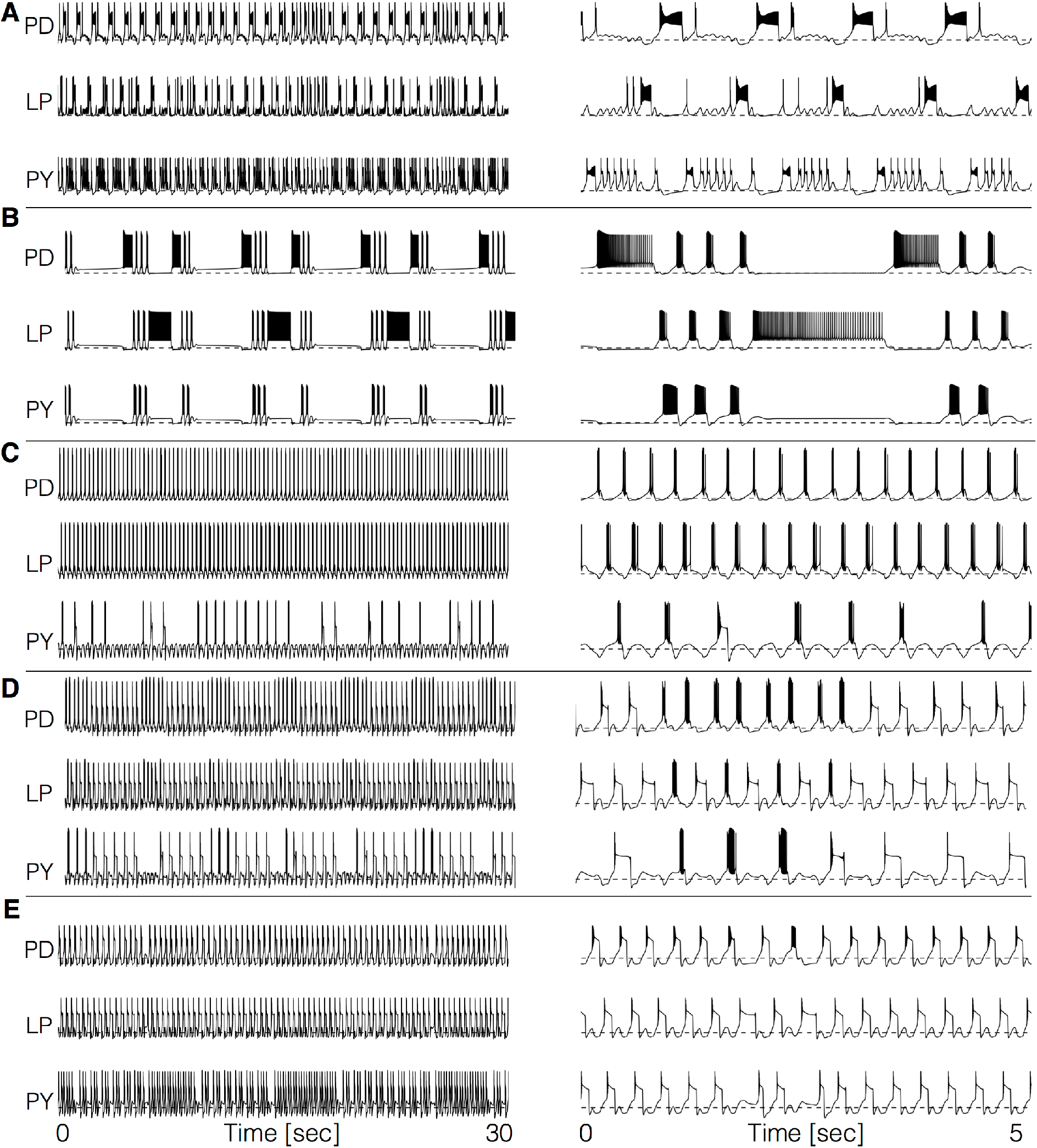
Dynamics of the circuits at high temperatures. The models become dysfunctional - or crash - at temperatures higher than 25°*C* and the do so in very different ways. The figure shows the membrane potential of five models **A-E**. The traces on the *left* show 30 seconds of data and the traces on the *right* show an expanded trace of 5 seconds. One hallmark of these regimes is the emergence of time scales much longer than that of the original rhythm.

### E. Dynamics of the circuits at high temperatures

For temperatures higher than a critical temperature, the biological network displays anomalous regimes characterized by the emergence of slow time scales and intermittency between what appear to be metastable states (Tang et al., 2012; Haddad and Marder, 2018). We explored how such regimes may arise in the models by direct inspection of the traces at different temperatures. We found that some models do not display irregular behavior but just become quiescent, or produce subthreshold oscillations, after a critical temperature. Many models do display irregular and seemingly aperiodic regimes. As expected, irregular regimes typically take place for values of the temperature near a transition between qualitatively different patterns of activity.

Figure 6 shows the activity of 5 models at one of their critical temperatures. The left panels show 30 seconds of data and the right panels show 5 seconds. Notice that in all cases there are timescales that are much larger than the control period of 1*sec*. The dynamics of the models in these regimes is daunting and its characterization is beyond the scope of this work. However, qualitative features of these states—such as the appearance of slow timescales or absence/presence of activity in one or more cells—are captured by these models. This, together with the multistable regimes shown before plus a source of noise, may provide a reasonable model to account for the irregular regimes in the biological network (Tang et al., 2012; Haddad and Marder, 2018).

### F. Dynamics of the currents at different temperatures

Neuronal activity is governed by currents that result from the precise activation and inactivation of ion channels with different kinetics. Experimental access to these quantities is limited since it is hard to measure currents individually without blocking all other currents, and thus changing the activity. Here, we employ the models to explore how these currents are differentially altered by temperature, and yet are able to remain balanced in such a way that the network activity speeds up while preserving its behavior and phase relationships.

We first compare the dynamics of the currents in two model *PD* neurons with different maximal conductances *G* and temperature sensitivities *Q*_10_ by direct inspection of the currents’ time series. Figure 7A shows the time series of each intrinsic current in *PD* at 10°*C* and 25°*C* for one example model. The corresponding membrane potential activity *V* is shown in the blue traces on top. The red dashed lines indicate the peak amplitude of the currents at control temperature (10°*C*) to visualize whether it grows or decreases. This is indicated by the boxes in each row (in colors: red grows, green decreases, white stays the same). In this example, the *Na* current increases its amplitude by more than two fold over the 15°*C* range. The *CaT* current also shows a strong dependence with temperature: despite the fact that *gCaT* increases with temperature, the amplitude of this current decreases to about half its control value at 10°*C*. Similar observations can be made for other currents (such as *I_A_* in Fig. 7A). The general case is that any current can increase or decrease depending on the dynamics of neuronal firing, the maximal conductances *G* and temperature sensitivities *Q*_10_. In addition, we found that whether any given current increases or decreases with temperature can be different across cells in the same model.

**FIG. 7.**
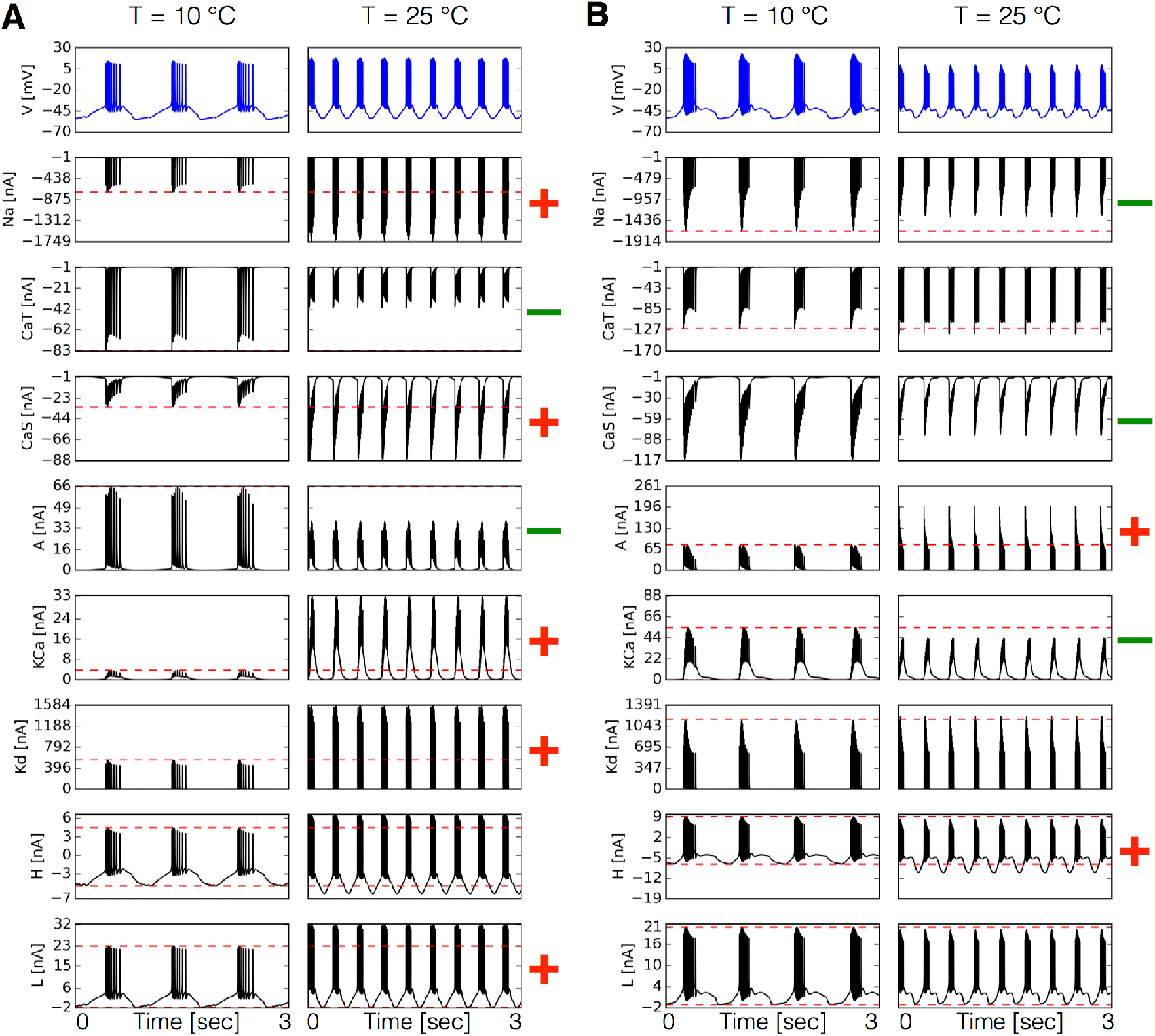
Dynamics of the currents at different temperatures. Currents of the *PD* cell at two temperatures for two different models **A-B**. Temperature increases the peak amplitudes of some currents and it decreases it for others, as indicated by the + and - symbols in red and green. The way these currents are modified can vary substantially across models.

The way currents are modified by temperature is different across models. Figure 7B shows the currents in the PD cell at two temperatures for another model. In this case, the *Na* current decreases by a small amount despite the fact that the maximal conductance and timescales of this current increase with temperature. Notice how the trends are different across models (Fig. 7A and 7B). The *KCa* current which in Fig. 7A grows by a factor six, is moderately decreased by temperature in Fig. 7B. Temperature not only affects the amplitudes of the currents but it also affects their waveforms in interesting ways. Because the total number of currents in the circuits is 31 it becomes cumbersome to compare them across temperatures (and models). For this reason, we computed and inspected their currentscapes (Alonso and Marder, 2019). The currentscapes use colors to show the percent contribution of each current to the total inward (or outward) current and are useful to display the dynamics of the currents in a compact fashion.

Figure 8 shows the currentscapes of each cell in one model at two different temperatures. The shares of each current of the total inward and outward currents are displayed in colors over time. The total inward and outward currents are represented by the filled black curves in logarithmic scale in the top and bottom. The currentscapes are noticeably different indicating that the currents contribute differentially across temperatures. For example, in the *PD* cell the *Leak* current contributes a visible share of the inward current both at the beginning and the end of the burst at 10°*C*, but at 25°*C* its contribution is mainly confined to the beginning of the burst. The *A* current is more evenly distributed during the burst at 10°*C* and at 25°*C* its contribution is visibly larger at the beginning of the burst. In all cells in this example model, temperature massively increases the contribution of the *KCa* current towards the end of the bursts. Figure 9 shows the same analysis for a second example. In this case also, the *KCa* currents increase their share at 25°*C* but is almost absent in the PD cell at 10°*C*. The *CaS* current increases with temperature in the PD cell, so temperature has the opposite effect on this current as it did in the example in Fig. 8. The *Leak* current contributes a large percent of the current both at the beginning and end of bursts at 10°*C*. At 25°*C* its contribution at the end of the burst becomes negligible while it still contributes significantly at the beginning of bursts. The contribution of the *CaT* current increases visibly in *PY*. In the third example in Figure 10 the contributions of the *KCa* current in the PD and *LP* cells are smaller and stay approximately constant. The contribution of the A current decreases in these cells as in previous examples, but because there is not an increment of *KCa*, the contribution of the synaptic currents—represented by the gray areas in the currentscapes—increases instead of decreasing. Together, these examples illustrate how a current can play different roles at different temperatures, and how diverse these mechanisms can be across individual solutions.

**FIG. 8.**
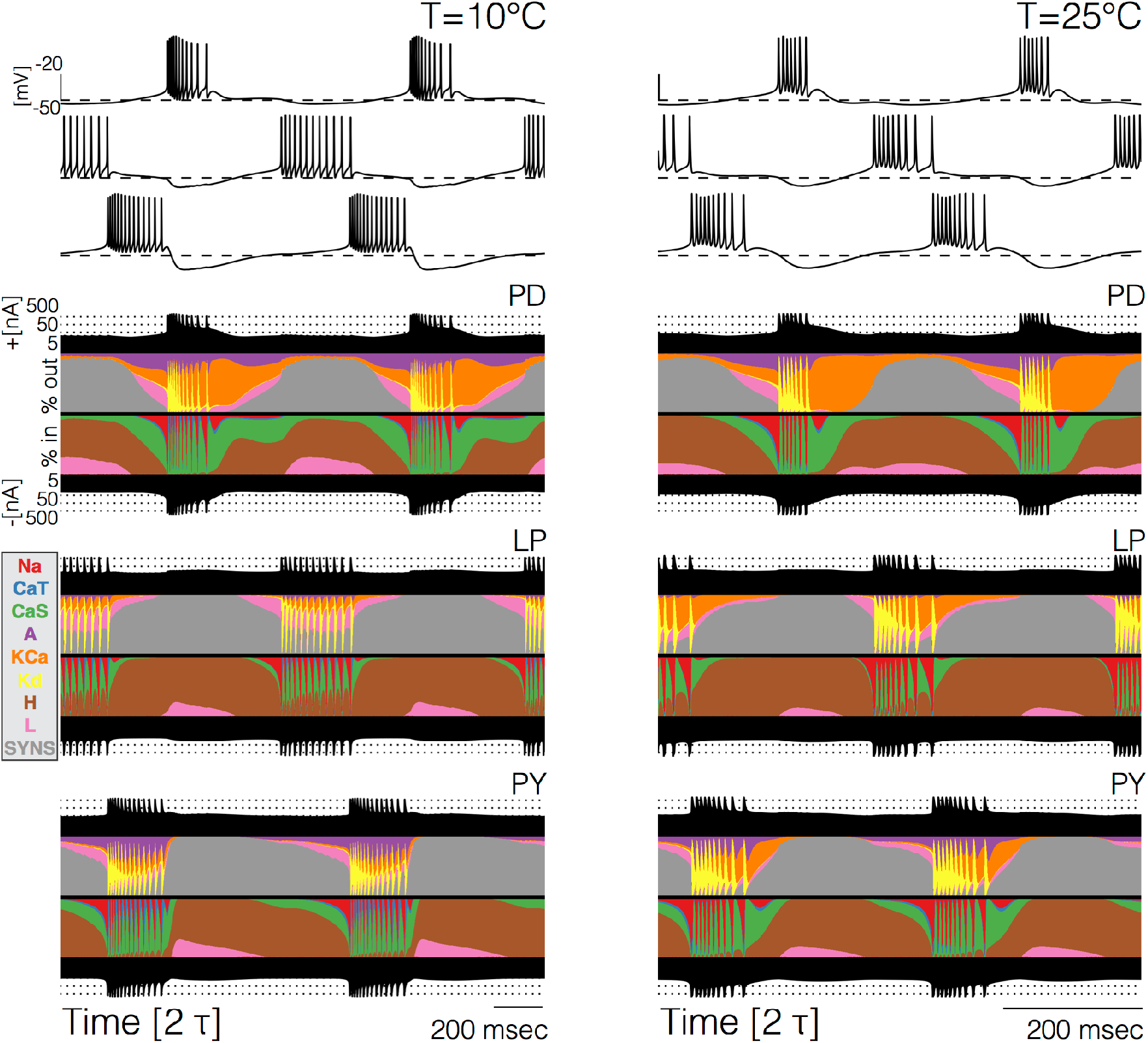
Currentscapes at different temperatures. The currentscapes use colors to display the percent contribution of each current type to the total inward and outward currents over time. The filled lines at the top and bottom indicate the total inward/outward currents in logarithmic scale. The figure shows the currentscapes for the three cells in the model at two temperatures. The panels show 2 periods *τ* of the oscillation.

**FIG. 9.**
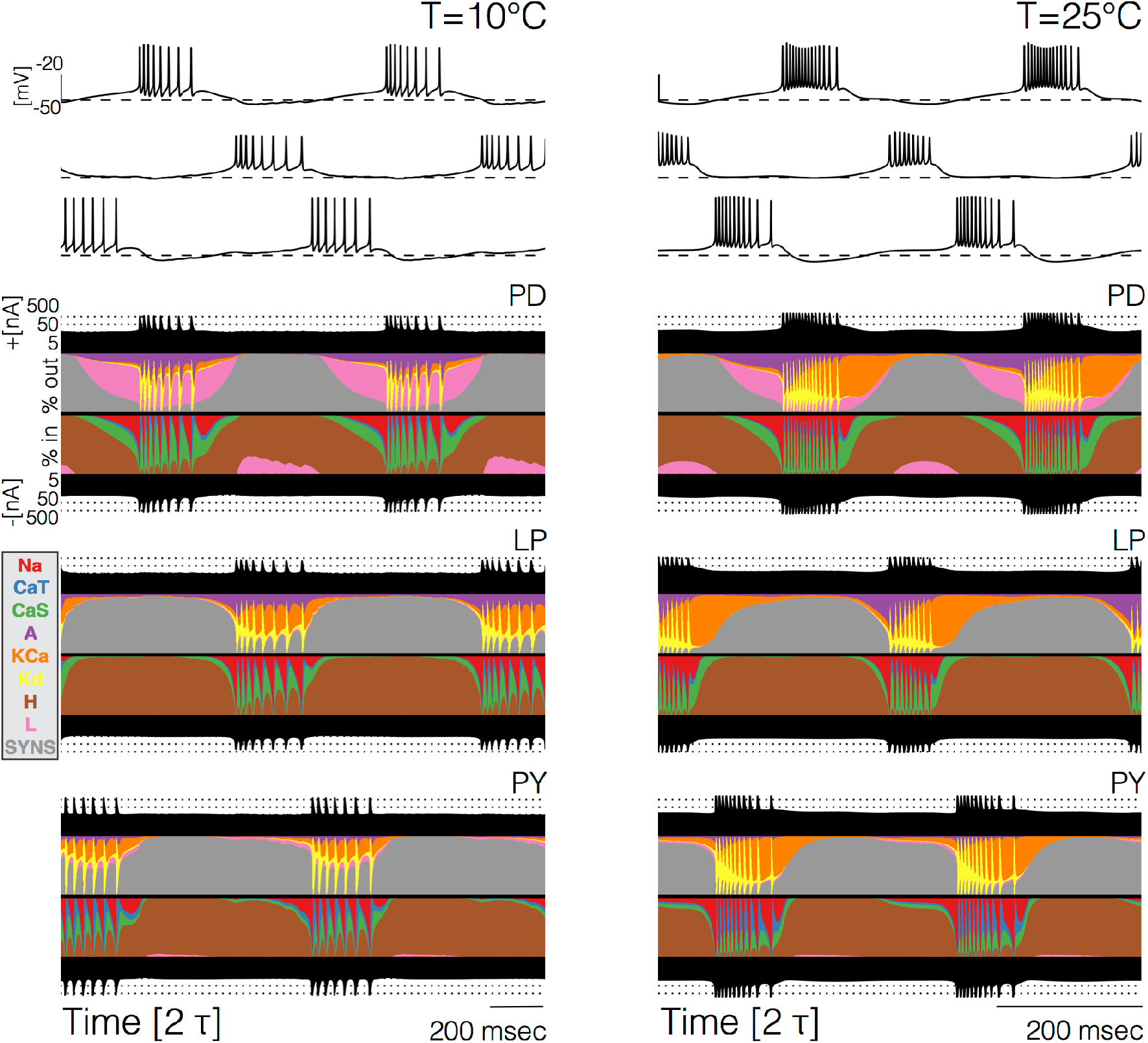
Currentscapes at different temperatures. Currentscape of each cell over two periods *τ* of the oscillation at two temperatures.

**FIG. 10.**
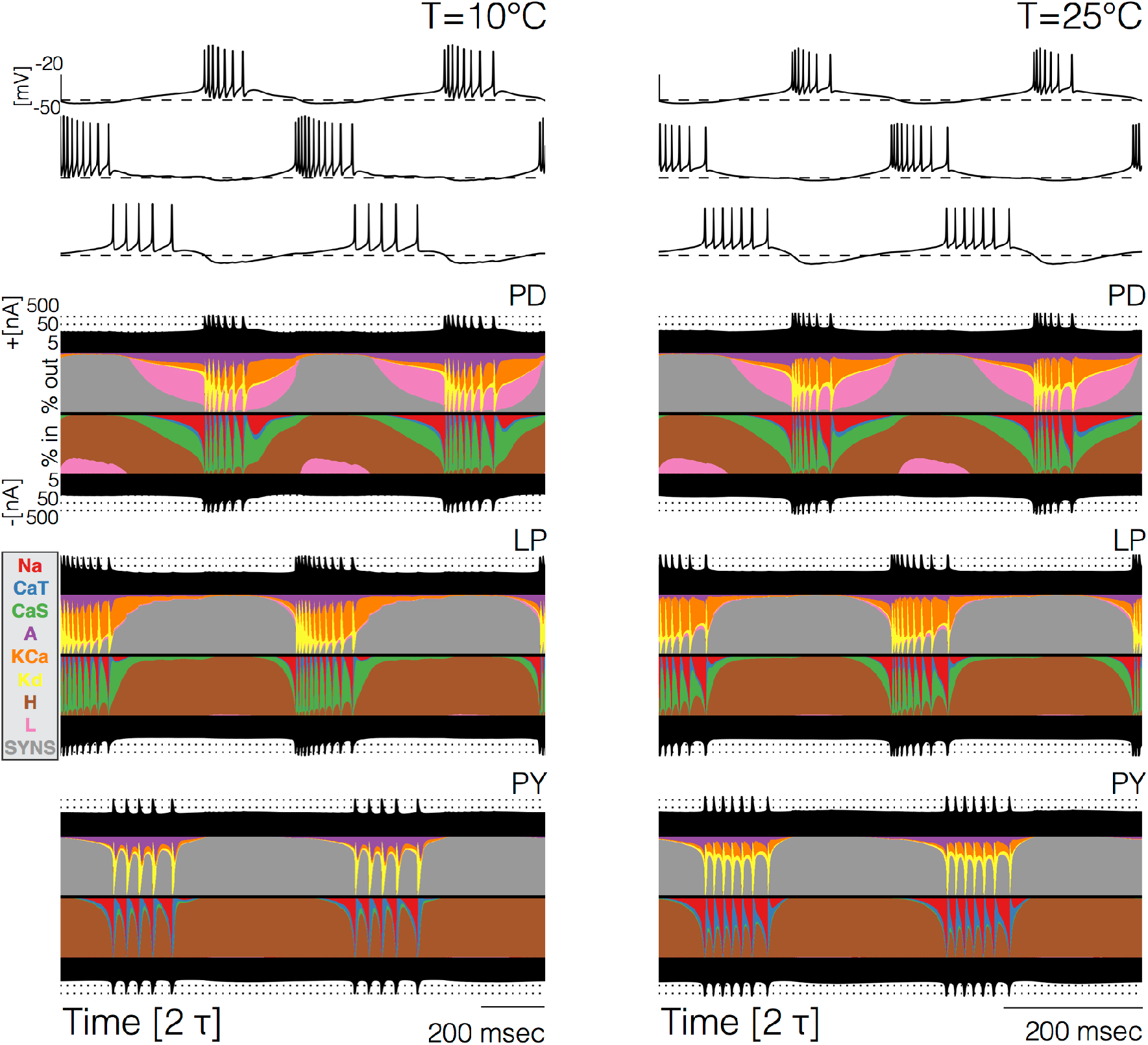
Currentscapes at different temperatures. Currentscape of each cell over two periods *τ* of the oscillation at two temperatures.

The currentscapes show that in all models and at all temperatures, there are periods of time where the activity is dominated by one or two currents, and periods of time where several currents contribute to it by similar shares. During spike production the activity is first driven by *I_N_a__* and later by *I_K_d__*. During these periods the total inward and outward currents differ by orders of magnitude, and are entirely dominated by one current type. Similarly, when the cells are most hyper-polarized the dominant currents are *I_H_* and the synaptic currents *I_Syn_*. In contrast, there are periods of time—such as the beginings and ends of bursts—where the total inward and outward currents are are composed of comparable contributions of several current types. It is during these periods—when multiple currents act together to change the membrane potential—where we observe the most dramatic changes with temperature. To show this we computed the currentscapes for the 20 milliseconds following the end of bursts in the PD cells of the models in Figures 8, 9 and 10.

Figure 11A shows how the currentscapes at burst termination change with temperature in the model in Fig. 8. In this case the inward currents show more changes than the outward currents: the contribution of the *CaT* current decreases with increasing temperatures and the contribution of *CaS* increases. The most salient changes in the outward currents are that *I_A_* decreases its contribution and the contribution of *I_KCa_* increases (orange areas). Differences across temperature are more visible for the model in Fig. 11B (also Fig. 9). In this model the *A* current also decreases its share as temperature increases but this effect is more pronounced than in Fig. 11A. The decrement in the share of *I_A_* is accompanied by a massive increase of the *KCa* current contribution. The contributions of the *Leak* and synaptic (*SYNS*) currents decrease for increasing temperatures. The inward currents also show visible differences. The contribution of the *Na* current increases and the contribution of the calcium currents *I_CaT_* and *I_CaS_* change visibly (their overall contribution appears to decrease, but their timing is also affected). The average contribution of the *H* current appears similar across temperatures but its dynamics is changed. It tends to increase its contribution after the end of the burst, and this increment becomes faster at higher temperatures. The third example in Fig. 11C (same model as in Figure 10) shows similarities and differences with the previous cases. The *A* current decreases its share but the contribution of the *KCa* current does not increase massively as in the previous cases. Instead, the contribution of the *Leak* current shows a large increment (pink areas). The inward currents also change with increasing temperatures: the contributions of the calcium currents *CaT* and *CaS* decrease, and the contribution of the *H* current increases. In total, Figure 11 shows that the ionic mechanisms that accomplish burst termination can be different across temperatures, and the way in which they change can be dramatically different across models. But importantly, in all cases there is a smooth transition between different relative fractions of currents. While all currents change their contributions with temperature, the most dramatic changes involve the relative roles of the *K*+ currents.

**FIG. 11.**
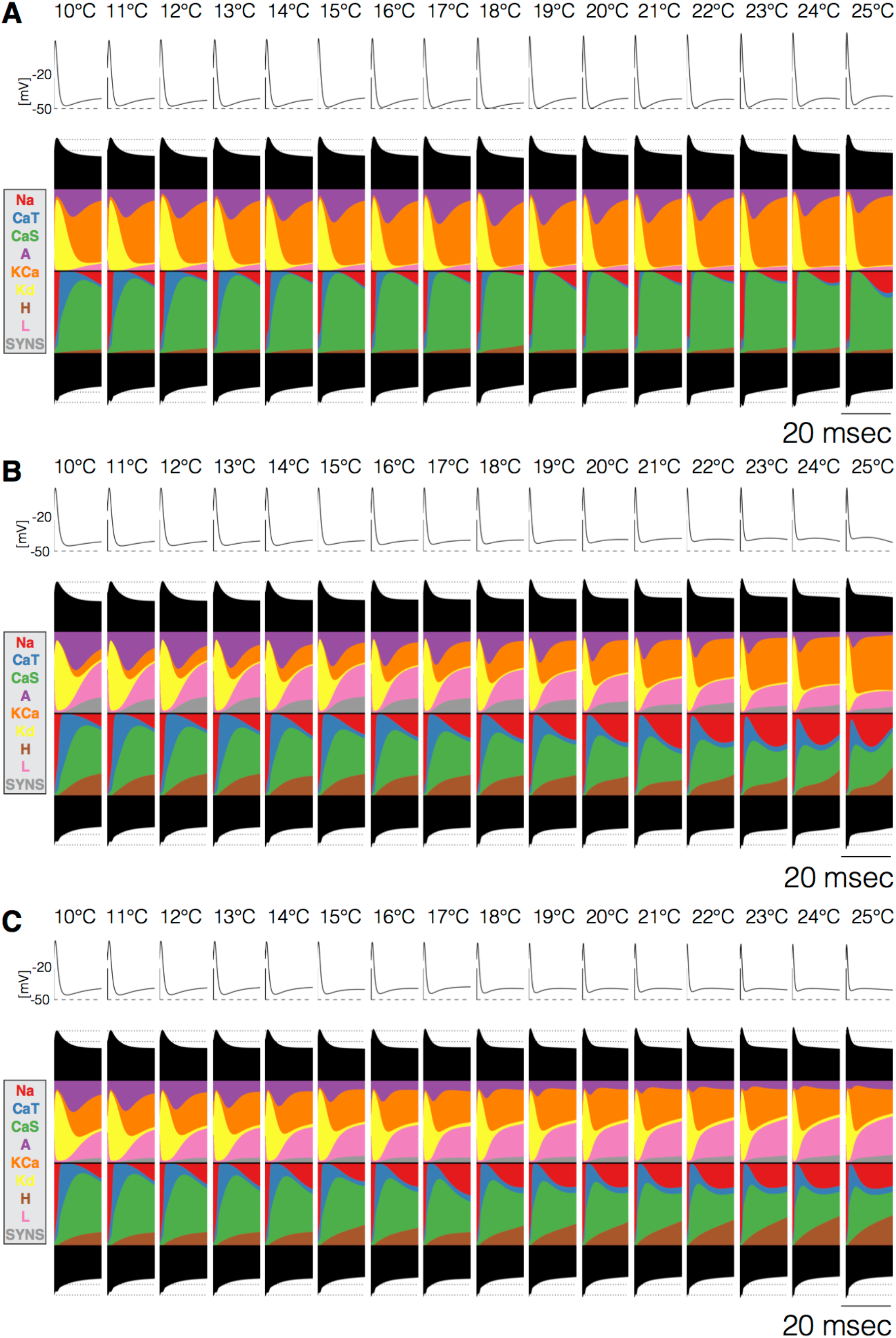
Currents’ contributions at the end of bursts across temperature. The figure shows the currentscapes of the PD cell for the 20 milliseconds following burst termination, and how these change over temperature. Panels **A-B-C** correspond to the models in Figs. 8, 9, and 10.

Figure 11 shows how the currentscapes at a specific instant change with temperature. To explore how the full currentscapes change over temperature we computed the distributions of current shares (Alonso and Marder, 2019). We computed the currentscapes at each value of temperature over the working range and inspected the time series of the currents’ shares, or equivalently, the widths of the colored regions in the currentscapes. We then proceeded in the same way as in Fig. 2 and computed the distributions of these time series at each temperature. Supplemental Figure S11A shows the time series of the *PD* cell in one model at two temperatures (10°*C* and 25°*C*) and their corresponding membrane potential distributions (left panels). The distributions are normalized so the y-axis indicates the probability of *V* (*p*(*V*)) in logarithmic scale. The quantity *p*(*V*) is proportional to the time the system spends at each voltage bin. We colored the traces using this probability as a color map. In this way, different portions of the cycle correspond to different colors: spikes appear in blue, sub-threshold activity in yellow, and brown and pink correspond to near-threshold activity. Notice that when the activity is faster this color code still parses the activity in a similar fashion. Supplemental Figure S11A and S11B shows the distributions of the current shares of all currents for 101 values of temperature between 10°*C* and 25°*C* (x-axis), for the models in Figures 8 and 9. In each panel, the y-axis corresponds to the share of each current to the total (inward or outward) and the colors indicate the portions of the cycle during which the current contributes by each amount. The distributions feature far more details than we are able to describe so we will only describe a few. For example in Fig. 11A, the *H* current in the PD cell contributes with less that 75% most of the time (yellow, hyper-polarized) at 10°*C* but its contribution increases almost linearly as temperature increases. The *Na* and *Kd* currents contributions span the full range (0% – 100%) but they contribute large amounts only during spikes (in blue), while most of the time (yellow, hyper-polarized) their contributions are small (≈ 5%). The *Leak* current contributes between 0% and 25% and while the distribution changes in a complicated manner, its average contribution does not appear to change noticeably. As a last example, notice that this is not the case in the *PY* where a clear decreasing trend can be observed for the *Leak* current. Figure 11B shows the same analysis for the model in Figure 10. These examples make the case that the way the currents contribute to activity can change quite dramatically with temperature and that the details of how these changes take place can be different across models, as well as across cells.

### G. Multiple mechanisms for temperature compensation

In these models, temperature increases the timescales and maximal conductances of all channels but its effect on the currents is far more diverse. As a way to characterize these changes we performed a series of simple measures on the time series of each current, and monitored these quantities over the working temperature range (10 – 25°*C*). For each current, we measured the peak amplitude (see Fig. 7 red dashed lines), the mean value over 30 seconds, and the mean share of the total (average along the y-axis in Fig. S11 for each temperature, or areas of the colored regions in the currentscapes) also over 30 seconds. Figure 12 shows examples of how the *CaT* currents in LP change over temperature, for many models indicated in different colors. The left subpanels show the peak amplitude, average, and average share of the *CaT* current over temperature, and the right sub-panels show a linear fit to each of those curves using the same color scheme. Note that in most cases the linear fit is a good approximation to the curve so this permits using the slope of the fits to compare trends across cells and models.

**FIG. 12.**
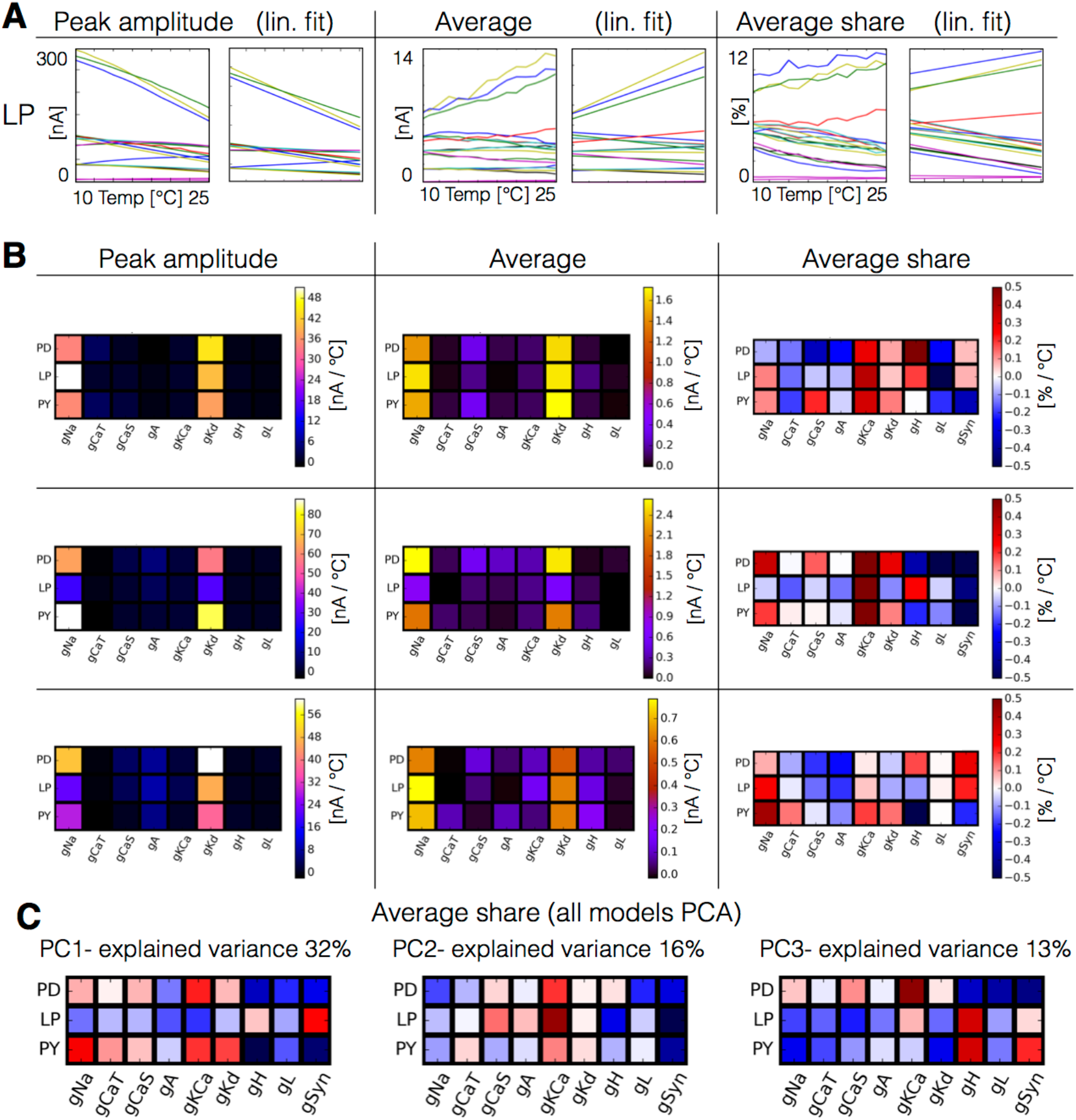
Summarizing changes in currents contributions with slopes. We quantified the effect of temperature on the currents by measuring their peak amplitudes, their average values and the average percent contributions to the total over 20 seconds. **A** The panels show each of these quantities for the *CaT* current in the LP cell, for 15 models indicated in different colors. The left sub-panels show the actual values and the right sub-panels show a linear fit to these trends. **B** Slopes of the linear fits in colors for each of these measures for the models in Figs.8-10 (top to bottom). The average share (right panels) shows more clearly that the way in which the currents change their contributions is diverse across models. The colors indicate whether a current increases its share (red) or decreases it (blue). **C** Principal component analysis of the average share trends. Color scale is the same as in **B**. The first three components capture ≈ 50% of the variance and show some correlated changes that occur frequently in the models studied here (*N* = 36).

These measures provide different information. The peak currents show the largest variations. The peak currents in one cell can either increase or decrease with temperature, and this trend can be different across cells despite their having the same *Q*_10_ values. The peak amplitude is not necessarily the most informative measure and it can be sensitive to large infrequent peaks such as those that occur in the synaptic currents when the spikes of two cells occur close in time (not shown). The average current already reveals different trends. The same currents whose peak amplitude decreased over temperature increase in the average. The third way in which we characterized these changes was by computing the average value of the shared contribution to the total. This corresponds to the width of each colored region in the currentscapes averaged over time. It is proportional to the total area of each colored region and therefore provides a measure of how the currentscapes change with temperature. This quantity behaves differently to the average current and is normalized to the interval [0,1].

Figure 12B shows the slopes of the fits of all current types in the three example models of Figs. 8, 9 and 10, for each of the quantities we monitored. In the first model (top row), the *Kd* current increases its peak amplitude by about 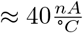 in all cells (less in PY). The *Na* peak current also increases in PD and PY but less than 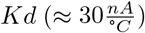 and it increases maximally in 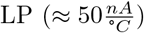. The peak amplitude of the calcium currents increases a small amount and the rest of the currents do not change their peak amplitudes. The second column shows the same analysis performed for the average value of the currents. This measure indicates similar trends as before showing growth of the *Na* and *Kd* currents, and it highlights that the average *CaS* currents increase more than the rest of the currents. Finally, the third column shows the trends of the average shares. In this case we used red for currents that increase their share and blue for currents that decrease it. The average contributions of currents can change by about ≈ 7% over the working temperature range and can increase or decrease. The second and third rows in Fig. 12B show the same analyses for the models in Figs. 9 and 10. The way the peak and average currents change with temperature appears to be similar across models, but if we consider the average contributions we see that each model achieves compensation by balancing its currents in different ways.

The mean current contributions or shares reveal more differences on how the currents become reorganized as temperature changes. The analyses in the third column of Fig. 12B can be compared directly with the currentscapes in Figs. 8, 9, and 10. The first row in Figure 12B, third column, shows in color whether a current increases or decreases its average contribution, for the model in Figure 8. In this model *KCa* grows in all cells. The contribution of the synaptic currents stays approximately constant in PD and LP but decreases in PY. The contributions of the *CaT* and *A* currents decrease in all cells by different amounts. The contribution of the *CaS* current changes differentially with temperature across cells: it decreases in *PD*, it increases in *PY*, and stays approximately constant in LP. All these trends can be seen by comparing the currentscapes at 10°*C* and 25°*C* as shown in Figure 8. In all cells in the second model (middle row, third column) temperature increases the average contribution of the *KCa* current and decreases the contribution of the synaptic currents. This is visible in the currentscapes as the *KCa* current takes more area at 25°*C* than at 10°*C* and reduces the average contribution of the synaptic current (area of the gray region in the currentscapes in Fig. 9). The third row in Fig. 12B shows this analysis for the model in Fig. 10. In this model the contribution of the *A* currents decreases in all cells. The contribution of *KCa* stays approximately constant in *PD* and *LP*, but it grows in PY. The contribution of the synaptic currents increases in *PD* and *LP*, but not in PY where it decreases. Overall, these three models achieve temperature compensation by completely different mechanisms.

There are multiple ways in which the currents can reorganize to produce a faster version of the activity while preserving its morphology. To gain intuition on how diverse these arrangements can be, we performed a PCA analysis across models on the trends described in Fig. 12A and 12B. We found that in the case of the peak amplitude, the first 2 components captured most of the variance (84%), which correspond to large changes in the *Na* and *Kd* currents. The same analysis performed with the average current also yields 2 components which capture most of the variance and are also very similar to the peak amplitude case (87%). The mean shares on the other hand require 6 components to explain a similar portion of the variance (84%), suggesting that this measure better highlights the different compensation mechanisms across models.

Figure 12C shows the first three Principal Components of the average shares’ slopes for all 36 models (right column 13B). The analysis suggests that there are certain currents that are more likely to change together. One example of this is indicated by the first component. If the *KCa* current increases (or decreases) its share, the synaptic current decreases (or increases) its share by a similar amount, and the *H* and *Leak* currents behave similarly to the synaptic current. The second component suggests that large increments in the *KCa* current in the *LP* neuron are usually correlated with changes in the *H* and synaptic currents in *PY* and anti-correlated with changes in the *Na* current. The third component shows that in both *LP* and *PY* an overall increment or decrement of the *Na* and *Kd* shares is anti-correlated with the *H* and synaptic currents.

### H. Response to perturbations at different temperatures

In all models the dynamics of the currents are different at different temperatures and their contributions become reorganized in complex ways. Therefore the role any given current plays at 10°*C* can be different at a different temperature (25°*C*). The combined activity of all the variables in the model—the dynamical attractor—is therefore expected to have different stability properties at different temperatures. This means that in the models, an extreme perturbation can have qualitatively different effects at different temperatures. The responses of different models to extreme perturbations such as partially or completely removing a current can be remarkably diverse at any temperature (Alonso and Marder, 2019). While this is expected due to the complexity of these models, the fact that these responses can be qualitatively different across temperatures is perhaps less clear. To shed light on this issue we performed simple perturbation protocols.

Figure 13 shows one perturbation protocol in two models (see models’ currents in Figs. 8 and 9). In Fig.13A we explored the interaction between temperature and removing the *A*-type *K*+ channel because Tang et al. (2010) measured the effects of temperature on *I_A_*. We performed the same perturbation at two temperatures: the models were simulated for 20 seconds in control conditions and then *I_A_* was completely removed for the next 20 seconds. Figure 13A (top) shows that removing the *A* current at 10°*C* has catastrophic consequences for the rhythm and the activity becomes completely irregular. When the same perturbation is performed at 25°*C* (bottom) the triphasic rhythm is almost normal except for the irregular bursting of *PY*. Figure 13B shows a similar pertubation protocol where we remove *KCa* in a different model (Fig. 9). This model displayed normal triphasic behavior at 10° (top) and when the *KCa* current was removed its activity slowed down but the triphasic rhythm was mantained. The same perturbation at 25°*C* results in quiescence. In this model, the *KCa* current is necessary for the activity at 25°*C* but not at 10°*C*.

**FIG. 13.**
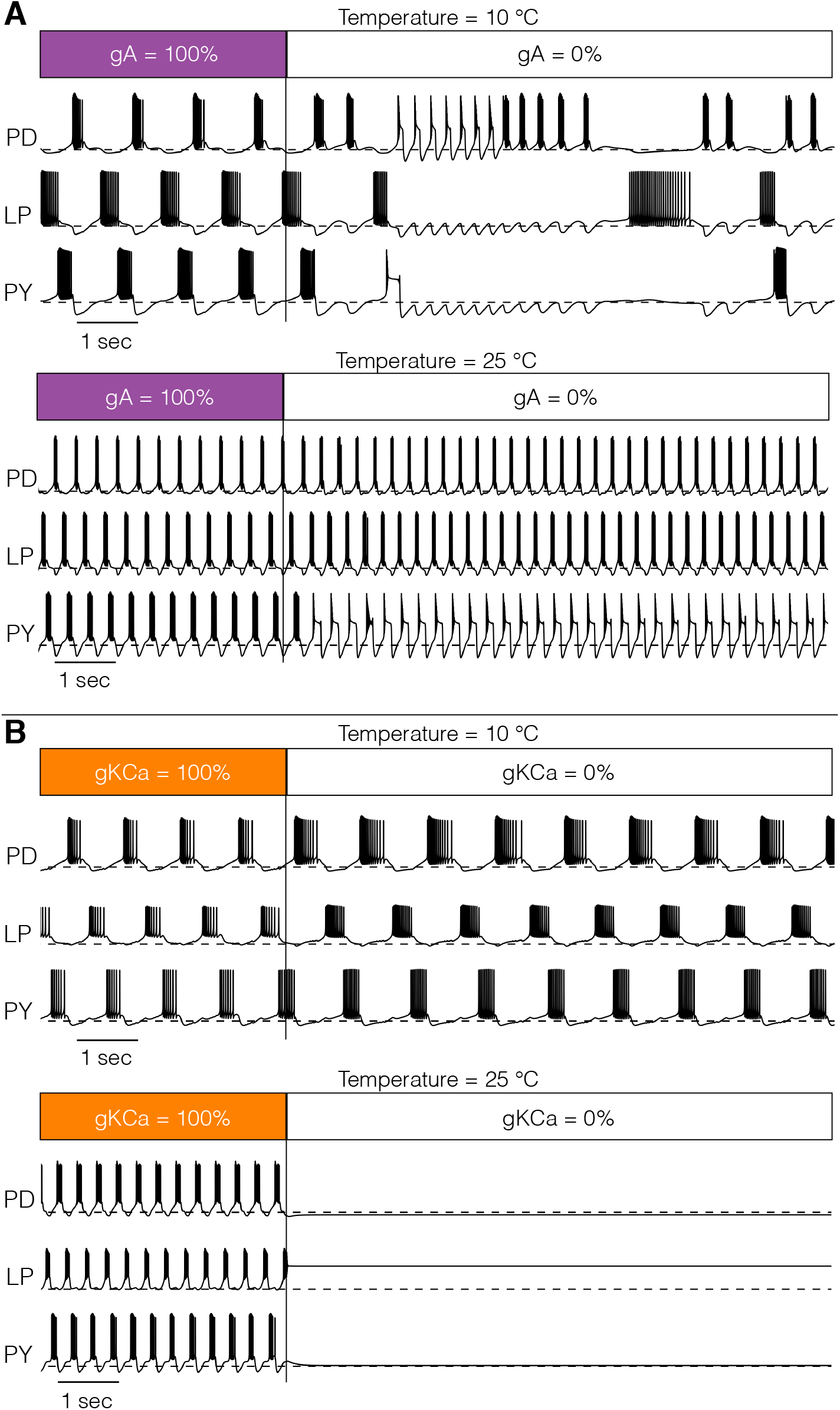
Response to perturbations at different temperatures. The response of the models to extreme perturbations can be different at different temperatures. The figure shows the responses of the models to the same perturbation at two different temperatures. **A** (top) Membrane potential over time at 10°*C*. The first 4 seconds correspond to the control condition with all currents intact. At *T* = 4*sec* we removed the A current. (bottom) Same perturbation performed at 25°*C*. **B** Same protocol as in **A** but removing *KCa* in a different model

To further characterize the differential response to perturbations at different temperatures, we performed a second assay where we gradually decreased a current from control (100%) to completely removed (0%) in five steps. We performed these simulations for all 36 models and all current types at 10°*C* and 25°*C* and compared their responses. In general, gradually removing a current can result in qualitatively different states depending on the temperature at which this perturbation is performed. The responses are so diverse that a coarse classification scheme is sufficient to highlight this observation. We classified the network activity based on the spiking patterns and waveforms of each cell, and also by their relative activities. The activities of each cell were classified as: regular bursting, irregular bursting, tonic spiking, irregular spiking, single spike bursting, quiescent and other, while the network activity was classified as triphasic or not triphasic. The classification scheme consists of a simple decision tree based on the statistics of spiking and is sufficient to tease apart the different regimes we observe at a coarse level (Methods).

Figure 14A shows the result of classifying the responses of one model to gradually decreasing a conductance in 5 steps, at two different temperatures (10°*C* and 25°*C*), for four conductances. We employed 5 × 4 *response grids* to summarize the effects of gradually removing a conductance at any given temperature. The rows in the grids correspond to different conductance removals, with the control condition (100%) on top, and the completely removed condition (0%) at the bottom. The first three columns correspond to each of the cells (PD-LP-PY) and the fourth column corresponds to the network (NET). The colors indicate the type of activity of each cell and the network at each condition as indicated by the labels in the figure. The top left panel in Figure 15A shows the responses of one model to removing the sodium current *I_Na_* at two temperatures. At 10°*C*, when *gNa* → 75%*gNa* the *LP* cell bursts irregularly and the activity of the network is *not triphasic*, but if the same perturbation is performed at 25°*C* the activity remains normal triphasic. Removing *KCa* in this model has little effect at 10°*C* as the activity remains triphasic. However, this current plays an important role at 25°*C* and the model becomes quiescent upon complete removal of *KCa*. The same observation that the responses to perturbation are different at each temperature holds regardless of the perturbation type. In the cases of *CaT* and *CaS* complete removal results in different non-triphasic activity at each temperature.

**FIG. 14.**
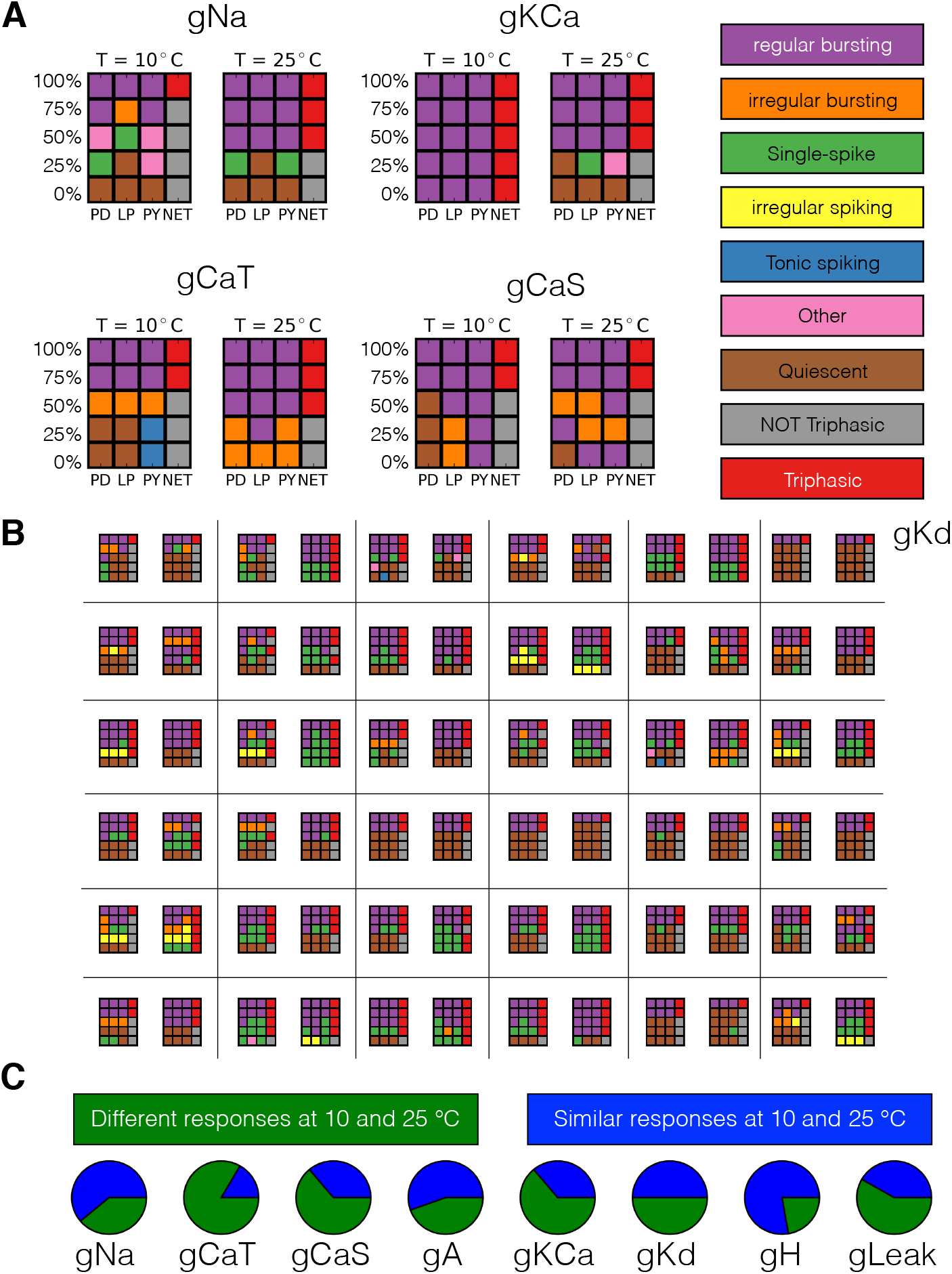
Classification of responses at different temperatures. The models’ responses to perturbations are extremely diverse and are in general different at different temperatures. We classified the responses of the models at two temperatures for 5 values of gradual decrements of each current. **A** Response to removal of *Na, A, CaT* and *CaS* for one model. **B** Response to removal of *Kd* at two temperatures for all models. C The pie charts show the number of models for which the responses to complete removal (0%) are classified as the same (green) or different (blue) at 10°*C* and 25°*C*.

Figure 14B shows the results of this analysis for all 36 models for the case of gradually removing *Kd* (labels are as in Fig. 14A and were dropped for clarity). The figure shows that the responses of the model are in general qualitatively different at different temperatures, and also that they are diverse across models. The fact that the models “break-down” in different ways is consistent with the observation that these compensation mechanisms are indeed diverse. In all models, the responses to extreme perturbation are temperature dependent for most perturbation types. In Figure 14C we quantify how many models respond differently to complete removal of a current at each temperature. This corresponds to comparing the bottom row of the response grids at each temperature, for each conductance type. In most cases, about half of the models respond to complete removal in different ways at each temperature. The exceptions are *CaT* for which most models respond in different ways, and *H* where complete removal results in equivalent states (quiescence in this case) at both temperatures. Overall our results indicate that the responses to perturbations can be qualitatively different at different temperatures.

## III. Discussion

Neurons characteristically have multiple ion channels that differ from each other in terms of their selectivity, voltage-dependence and time courses of activation and inactivation. For example, it is not uncommon that neurons might display 12 or more different voltage and time-dependent currents, including multiple classes of potassium and calcium currents. Undoubtedly, many of these channels are targeted to specific sites on neurons, but nonetheless they all will contribute to the overall electrical activity of neurons and the networks in which they are found. One of the potential advantages of this multiplicity of channels is that they may confer resilience to perturbations of many kinds (Drion et al., 2015). In this paper, we explored some ways in which multiple ionic currents help confer temperature robustness to neurons and a small network.

Temperature affects the properties of ion channels differentially and yet many animals live successfully over wide temperature ranges. The North Atlantic lobsters and crabs routinely experience more than 25°*C* temperature swings, and may see 8 – 10°*C* changes within a short time period.

The temperature sensitivities of ion channels are dictated by their molecular structures, and for this reason it may be reasonable to assume that the temperature sensitivity of a given channel is similar across cells and individuals, and throughout the lifetime of the animal. This raises the question of whether it is possible for one set of *Q*_10_ values to function appropriately over wide ranges of conductance densities. Alternatively one may consider molecular processes which could alter channel function. For example, RNA editing can change the temperature sensitivities of specific channels to facilitate adaptation (Garrett and Rosenthal, 2012; Rosenthal, 2015). A recent study in *Drosophila* showed that acute temperature perturbations result in changes in RNA editing levels (Rieder et al., 2015). Therefore, it is also reasonable to explore the possibility that the *Q*_10_ values can be different across individuals and change over time.

Nervous systems must have mechanisms that allow them to rapidly and reliably deal with acute changes in temperature. We provide simulations that suggest that some sets of ion channels are able to deal with changes in temperature of this sort by smoothly changing the mechanisms by which neuronal activity is produced. This can be seen most dramatically in Figure 11 which shows that the role of the three different *K*+ currents in spike repolarization gradually changes as the temperature increases. While this is true in all three examples shown in Figure 11, the details by which the currents trade-off their role in spike repolarization are different, and depend on the differences in their starting conductance values. This smooth transition can occur precisely because these three *K*+ depend on voltage and time differently and also are affected differently by temperature. Our speculation is that this kind of smooth transition is one of the reasons that neurons have so many ion channels with overlapping functions.

Transitions in the relative importance of different currents are also seen as a function of changes in membrane potential (Golowasch et al., 1992) and as a function of spike broadening in a repetitively firing neuron (Ma and Koester, 1996). So it is likely that the smooth transition in the relative roles of multiple conductances in neuronal dynamics is a general feature of all neurons with complex dynamics and many kinds of channels. This illustrates the benefit for a neuron to have multiple channels with overlapping functions.

The same principle is likely to hold when we approach network dynamics. The phase relationships of the pyloric rhythm are maintained over a considerable temperature range (Tang et al., 2010; Soofi et al., 2014). Likewise, the pyloric phase relationships are also maintained over a considerable frequency range (Hamood et al., 2015). In this latter case, it has been argued that the phase compensation depends on the conjoined action of a number of different cellular mechanisms, including synaptic depression (Manor et al., 1997; Hooper, 1997), and the activation and inactivation of *I_A_* and *I_H_* (Nadim et al., 1999; Nadim and Manor, 2000; Tang et al., 2010; Harris-Warrick et al., 1995). So, here the principle also holds: resilience and robust function may require smoothly moving between a variety of different cellular mechanisms.

In this work we employed biophysically detailed models of the pyloric network to explore how temperature compensation can be achieved in small neural circuits. We extended a model of the pyloric network developed previously to include temperature sensitivity and produced multiple temperature compensated models that capture much of the phenomenology reported in experiments (Tang et al., 2010, 2012; Soofi et al., 2014). One of the most significant results of this paper is that temperature compensation can take place in multiple different ways. In these models, both the membrane potential and the spiking patterns are affected by temperature changes and these changes appear to be different in every model we inspected. Across the models, different values of the maximal conductances *G* and temperature sensitivities Qι_0_ result in consistent differences in the duty cycle distributions. Measurements of the values of the maximal conductances in the STG show large variability across individual animals (Goaillard et al., 2009; Schulz et al., 2006, 2007) so our expectation is that the duty cycle distributions of the biological cells will also display intricate dependencies with temperature, and that these distributions will be different across individuals.

Temperature is not the only perturbation that crabs and lobsters experience. As with temperature, the responses of the STG to changes in pH are diverse across individuals (Haley et al., 2018), again consistent with the large amount of animal-to-animal variability in the expression of ion channels (Schulz et al., 2006, 2007; Temporal et al., 2014; Tran et al., 2019). It is therefore reasonable to assume that the responses to a global perturbation are diverse across individuals because different compositions of channel densities—which produce similar pyloric rhythms—are differentially resilient to any given perturbation. By the same token, as currents change as a function of temperatures we expect that a global perturbation of the same individual may have qualitatively different effects at different temperatures, as is illustrated in Figure 13. The interaction between temperature and a second perturbation are the subject of recent experimental studies (Haddad and Marder, 2018; Ratliff et al., 2018). These studies are consistent with the interpretation that different current configurations have different stability properties and that temperature changes these configurations.

In this work we used a genetic algorithm (Alonso and Marder, 2019) to find temperature-compensated networks. It is important to reiterate that temperature-compensated neurons and networks are difficult to find doing random searches (Caplan et al., 2014). Therefore, despite the significant animal-to-animal differences in conductance densities seen across the population, randomly sampling conductance values is unlikely to result in successful solutions for individual animals. This argues that the biological mechanisms that give rise to successful temperature compensation are likely to result from rules by which correlated values of conductances are produced, such as those found in homeostatic models (O’Leary and Marder, 2016; O’Leary et al., 2013).

Taken at face value, models suggest that conductance densities drift throughout the life of an animal as a result of homeostatic processes (LeMasson et al., 1993; Liu et al., 1998; Golowasch et al., 1999b; O’Leary et al., 2014). Conductance densities can also change in an activity dependent manner as a result of perturbations (Turrigiano et al., 1994; Golowasch et al., 1999a; Santin and Schulz, 2019; Golowasch, 2019). Understanding how temperature compensation can be compatible with homeostatic and adaptation mechanisms remains an open theoretical and experimental question.

Although the models used here are semi-realistic they are far from incorporating the complexity of actual biological neurons and the pyloric circuit. Therefore, we do not expect that specific observations derived from our cohort of models—such as the co-regulation of some current pairs being more likely than other pairs—have predictive value in detail. Nonetheless, we do expect that the general principles highlighted here will hold in biological systems. In particular, the observation that the dynamics of the currents are reorganized in non-trivial ways in response to changes in temperature is likely to be a common finding in biological systems.

## IV. Methods

### A. The model

The activity of the cells was modeled using single-compartment models similar to those described previously (Turrigiano et al., 1995; Liu et al., 1998; Goldman et al., 2001; Alonso and Marder, 2019). Each neuron has a sodium current, *I_Na_*; transient and slow calcium currents, *I_CaT_* and *I_CaS_*; a transient potassium current, *I_A_*; a calcium-dependent potassium current, *I_KCa_*; a delayed rectifier potassium current, *I_Kd_*; a hyperpolarization-activated inward current, *I_H_*; and a leak current *I_leak_*. The number of state variables per cell is 13. The units for voltage are *mV*, the conductances are expressed in *μS* and currents in *nA*. The pyloric network consists of 3 compartments with the same ion channel types but different conductance densities. The interactions in the network consist of 7 chemical synapses and are similar to Prinz et al. (2004). The synaptic current is given by *I_s_* = *g_s_S*(*V_post_*–*E_s_*), where *g_s_* is the synapse strength, *V_post_* is the membrane potential of the postsynaptic neuron and *E_s_* is the reversal potential of the synapse. The activation of a synapse *s*(*t*) is given by

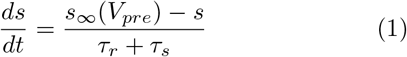

with,

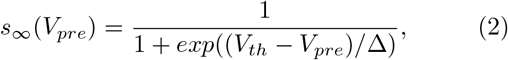

and

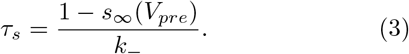

These equations are identical to Prinz et al. (2004) except for the inclusion of a bound for the timescale of activation *τ_r_* = 20*msec* (we want to avoid the case that as *s_∞_* → 1 then *τ_s_* → 0 and *ṡ* → *∞*). All other parameters (except *g_s_*) are identical to Prinz et al. (2004). Following Prinz et al. (2004) we set *E_s_* = −70*mV* and 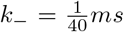 for glutamatergic synapses, and *E_s_* = −80*mV* and 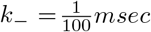 for cholinergic synapses. We set *V_th_* = −35*mV* and Δ = 5*mV* for both synapse types.

Temperature effects were included in this model as done previously by Caplan et al. (2014) and O’Leary and Marder (2016). Temperature dependence was introduced in the time constants of the channel-gating variables *τ_m_i__* and *τ_h_i__*, the maximal conductances *g_i_*, the time constants of calcium buffering *τ_Ca_*, and the maximal conductances and time constants of the synapses. This was done by replacing all conductances *g_i_* by *R_i_*(*T*)*g_i_* and all time scales *τ_i_* by *R_i_*(*T*)^−1^*τ_i_* where

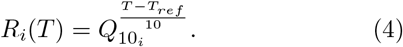

Here *T* is the temperature, *T_ref_* = 10° is the reference temperature and *Q*_10_*i*__ is defined as the fold change per 10°*C* from the reference temperature (*i* indicates the process type). Finally, the calcium reversal potential *E_Ca_* depends on temperature through the Nernst equation.

The total dimension of the model is 46 = 3 × 13 + 7 with 13 state variables per cell, and 7 variables for the state of the synapses. In this work some parameters were considered fixed while others were allowed to take values in a range. The number of parameters we varied per cell is 27 = 9 × 3: the 8 maximal conductances plus the calcium time constant *τ_Ca_*. The temperature sensitivities were assumed to be the same for all cells and amount to 24 additional parameters. We need to specify 8 *Q*_10_ values for the intrinsic maximal conductances (one per channel type), plus 12 *Q*_10_ values for the time scales of each intrinsic process (not all currents inactivate). The number of *Q*_10_ parameters for the synapses is 4: 2 for the maximal conductances (of each synapse type), and 2 for the timescales of activation. The models were simulated using a Runge-Kutta order 4 (RK4) method with a time step of *dt* = 0.05*msec* (Press et al., 1988). We used the same set of initial conditions for all simulations in this work *V* = −51*mV*, *m_i_*, *h_i_* = 0 and [*Ca*^+2^] = 5*μM*.

### B. Finding parameters

Each model is specified by the maximal conductances *G* and calcium time constants of each cell (9 × 3 parameters), and the temperature sensitivities *Q*_10_ of each process (24 parameters). We recently introduced a function that upon minimization, results in values of the maximal conductances for which the activity of a single compartment corresponds to periodic bursting with a target frequency (*f_tg_*) and duty cycle (*dc_tg_*). The function uses thresholds to obtain temporal information of the waveform, such as spike times, and then uses them to assign a score or error that measures how close the solutions are to a target activity. This function is described in detail in Alonso and Marder (2019) and was used here to find temperature compensated networks.

For any given set of conductances we simulated the model for 20 seconds and dropped the first 10 seconds to minimize the effects of transient activity. We then computed the average (<>) burst frequency < *f_b_* >, the average duty cycle < *dc* >, the number of crossings with a slow wave threshold #_*sw*_ = −50*mV*, the number of bursts #_*b*_, and the average lags between bursts < Δ_*PD-LP*_ > and < Δ_*LP-PY*_ >. To discard unstable solutions we checked if the standard deviation of the burst frequency and duty cycle distributions was small; a solution was discarded if *std*({*f_b_*}) ≥< *f_b_* > ×0.1 or *std*({*dc*}) ≥< *dc* > ×0.2. If a solution is not discarded we can use these quantities to measure how close it is to a target behavior,

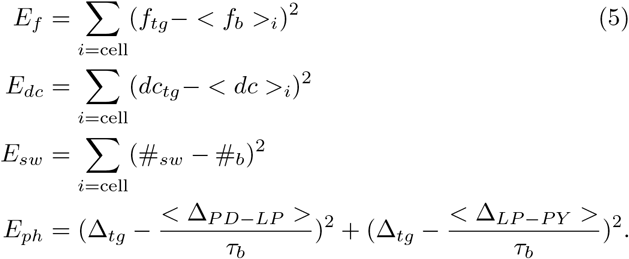

Here *E_f_* measures the mismatch of the bursting frequency of each cell with a target frequency *f_tg_* and *E_dc_* accounts for the duty cycle. *E_sw_* measures the difference between the number of bursts and the number of crossings with the slow wave threshold *t_sw_* = −50*mV* (we ask that #_*sw*_ = #_*b*_). *E_ph_* compares the lags between bursts (in units of the bursting period 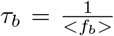) to a target lag Δ_*tg*_. These measures are discussed in more detail in Alonso and Marder (2019).

Let ***G*** denote a set of maximal conductances (and *τ_Ca_*), we can then define an objective function

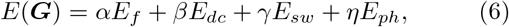

where the weights (*α,β,γ,η*) define how the different sources of penalties are pondered. In this work we used *α* = 10, *β* = 1000, *γ* = 1, *η* =10. The penalties *E_i_* were calculated using *T* = 10 secs with *dt* = 0.05 msecs. The target control behavior was defined as having all cells bursting with %20 duty cycle for *PD* (*dc_tgpD_* = 0.2) and 25% for the LP and PY cells. The lag between bursts was targeted to be Δ_*tg*_ ≈ 0.3.

Minimization of the objective function produces sets of maximal conductances *G* for which the resulting circuit activity mimics the pyloric rhythm. The minimization was performed over a search space of allowed values listed here: for each cell we searched for *g_Na_* ∈ [0,10^3^] ([*μS*]), *g_CaT_* ∈ [0, 10^2^], *g_CaS_* ∈ [0,10^2^], *g_A_* ∈ [0, 10^2^], *g_KCa_* ∈ [0,10^3^], *g_Kd_* ∈ [0,10^2^], *g_H_* ∈ [0, 10^2^], *g_L_* ∈ [0,10], *τ_Ca_* ∈ [0, 2 × 10^3^] ([msecs]). All synaptic conductances were searched in the range *g_syn_* ∈ [0, 5 × 10^-2^] ([*μS*]). We discretized our search space by taking 10^3^ equally spaced values for each parameter. We minimized the objective function using a custom genetic algorithm (Holland, 1992).

Temperature compensation is achieved in our models by searching sets of ***Q***_10_ values that produce the target pyloric rhythm at each temperature. As Caplan et al. (2014) and O’Leary and Marder (2016) we searched for values of *Q*_10_*i*__ between 1 and 2 for conductances and between 1 and 4 for activation/inactivation time scales. We built a new objective function by evaluating the previous objective function (6) over a set of control temperatures *T_i_* taken at 15°*C*, 20°*C*, 25°*C* and 35°*C*,

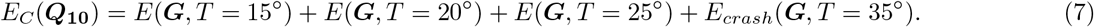

Here, *E*(***G**,T_i_*) is the score of the solutions of the set of conductances ***G*** at temperature *T_i_* and *E_crash_* is the total number of spikes in all three cells. This last term is introduced to enforce that when the temperature is close to 35°*C* the network stops working as in experiments. Evaluation of the objective functions (6) and (7) requires that the model is simulated for a number of seconds and this is the part of the procedure that requires most computing power. Longer simulations will provide better estimations for the burst frequency and duty cycle of the solutions, but it will linearly increase the time it takes to evaluate them. If the simulations are shorter, evaluations of the objective function are faster but its minimization may be more difficult due to transient behaviors and its minima may not correspond to stable pyloric rhythms.

Because we only target the activity at 3 temperature values, it is not guaranteed that the models will behave properly for temperatures in between. We found that if the system meets the target activity at the control temperatures, in the vast majority of the cases, the models also displayed the correct activity for temperatures in between, but there were exceptions. This means that even if a solution achieves low scores, we are still required to screen the temperature values in between to make sure it is indeed temperature compensated over the full range. Increasing the number of control temperatures makes the estimation procedure slower and we met a good balance in 4 control temperatures plus screening as described.

### C. Membrane potential and current shares distributions

The membrane potential distributions and current shares distributions were computed for 101 values of temperature between 10°*C* – 25°*C*. For each temperature, the models were simulated from identical initial conditions for 30 seconds with high numerical resolution (*dt* = 0.001). The distributions were computed using the last 10 seconds of each simulation. The number of samples of the distributions at each temperature is 5 × 10^5^.

### D. Classification scheme

We classified the activity of each cell into several categories by inspecting their spiking patterns and features of their waveforms. For this we first compute the spike times and their ISI distributions. If the coefficient of variation *CV_ISI_* < 0.1 we declare the activity as spiking. We then measure the lag *δ* between the spike onset and the crossing times with a slow wave threshold at −50*mV*. If this *δ* < 20*msec* we label the trace as *tonic spiking* and else, we label it as *single-spike bursting*. If *CV_ISI_* > 0.1 then we ask if the cell is bursting. For this we group spikes into bursts using a temporal threshold of 200*msec* and compute the distributions of duty cycle dc and instantaneous burst frequency *f_b_*. If *CV_dc_* < 1, *CV_fb_* < 0.1 and *mean*(*dc*) > 0 we label the trace as *regular bursting*. If *mean*(*dc*) > 0 and either *CV_dc_* > 1 or *CV_fb_* > 0.1 we label the trace as *irregular bursting*. If *mean*(*dc*) = 0 we label the trace as *irregular spiker*. If none of these conditions were met we label the trace as *other* and finally, if the total number of ISI values is not greater than 1 we label the trace as *quiescent*. The network state was classified as *triphasic* or *not triphasic*. We declared the network (NET) activity as *triphasic* if the frequency of the cells was similar and they fired in the right order, and not triphasic elsewhere.

### E. Parameters used in this study

Each model was assigned a 6 character name. Here we list which model was used in each figure. Code to simulate the network and the parameters of each model are supplemental to this work.

Figure 1: 4X6W66. Figure 2B, in lexicographical order, G7P2WE, 4X6W66, MAXTTP, PCR6RP, Q8FBN1, B7SDOL. Figure 3, PLKTEM. Figure 4, all models. Figure 5, OSHNMD, H5AIZR. Figure 6 (top to bottom), J9U8SQ, G81PUL, G7EOZ8, SAXT8Y, KNMLBC. Figure 7, 4X6W66, SJR46Y. Figure 8, WWZ3CN. Figure 9, FGFKQS. Figure 10, 71G6LA. Figures 11 and 12, WWZ3CN, FGFKQS, 71G6LA. Figure 13, G7EOZ8, FGFKQS. Figure 14, 4X6W66.

## Author contributions

LA: design of study, carried out simulations, prepared figures, wrote and edited the manuscript. EM: design of study, wrote and edited the manuscript.

## Funding acknowledgments

Research supported by MH046742 and R35 NS097343 (EM), T32 NS07292 (LA) and Swartz Foundation 2017 (LA).

## V. Supplemental Figures

**FIG. 1 Supplemental.**
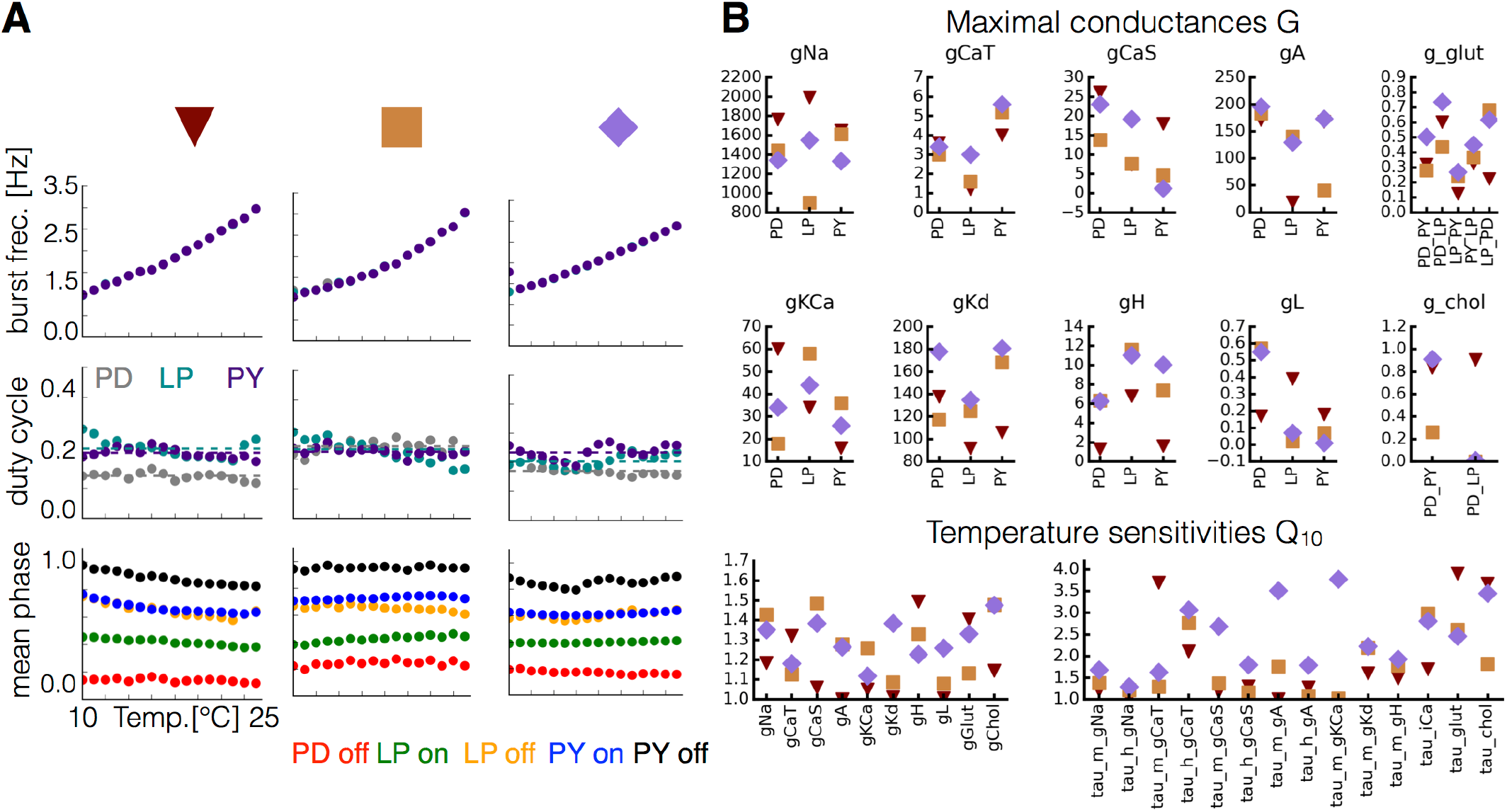
Equivalent phenotype for different sets of maximal conductances and temperature sensitivities. For each set of maximal conductances we studied, it was possible to find multiple sets of *Q*_10_ that yielded temperature compensated solutions. **A** Burst frequency (top), duty cycle (middle), and phases (bottom) over the working temperature range for three different models (indicated in colors). **B** Values of the maximal conductances (in *nS*) and *Q_10_* for each model.

**FIG. 11 Supplemental.**
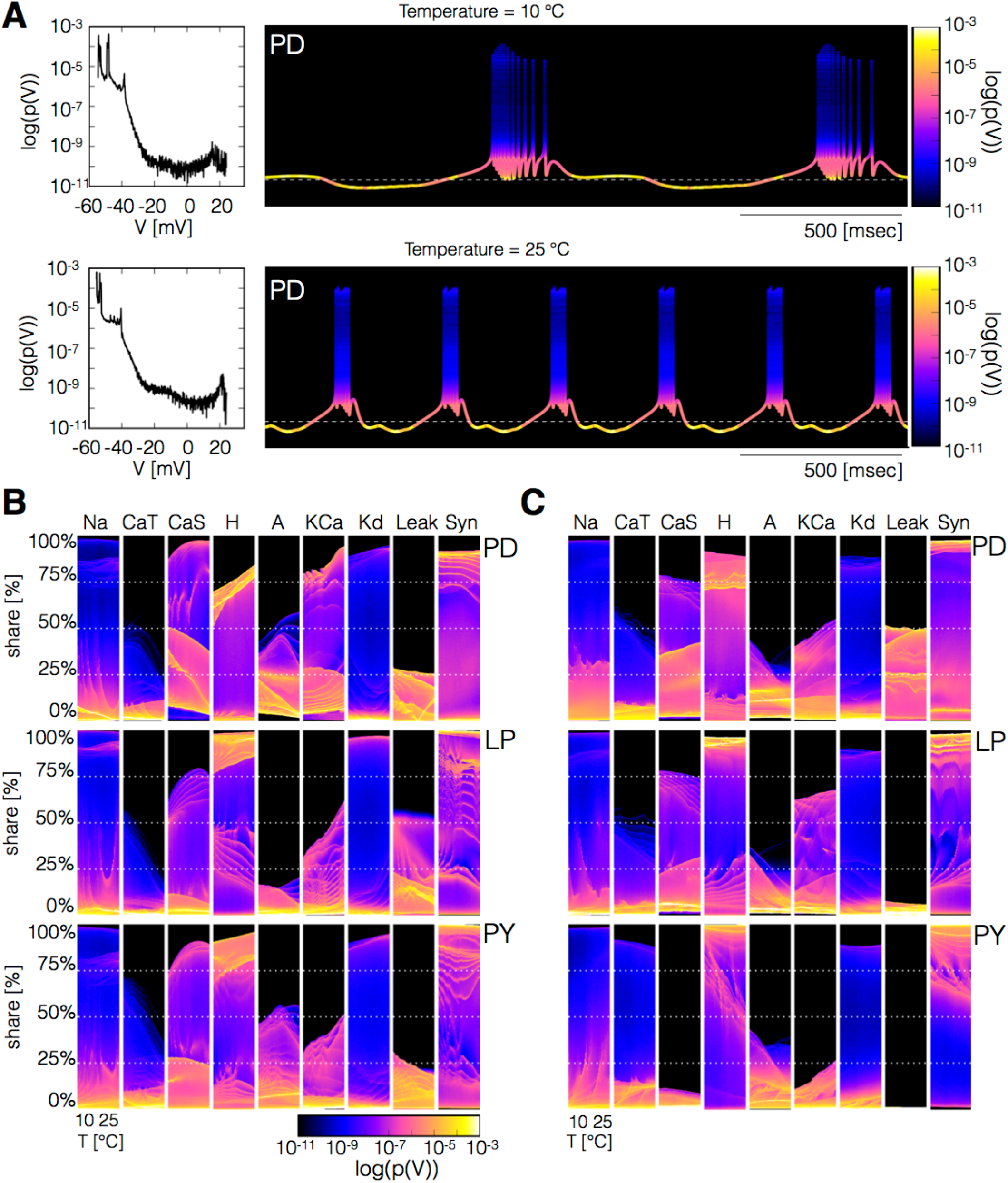
Distributions of current shares over temperature. **A** Traces and membrane potential distribution at two temperatures (top: 10°*C*, bottom: 25°*C*). Traces were colored using *log*(*p*(*V*)) as a color scale. In this way, different portions of the cycle appear in different colors: blue for spikes, yellow for sub-threshold activity and brown and pink for near threshold activity. **B-C** The panels show the percent contribution of each current to the total inward (and outward) currents, over the working temperature range (x-axis, 10°*C* – 25°*C*)). The colors indicate the portions of the cycle at which each current contributes by each amount (y-axis).

## Notes

#### Summary of Updates

Abstract was updated. Figure 13 was relabeled.

## References

Abrams, T. W. and Pearson, K. G. (1982). Effects of temperature on identified central neurons that control jumping in the grasshopper. Journal of Neuroscience, 2(11):1538–1553.

Alonso, L. M. and Marder, E. (2019). Visualization of currents in neural models with similar behavior and different conductance densities. eLife, 8:e42722.

Bucher, D., Taylor, A. L., and Marder, E. (2006). Central pattern generating neurons simultaneously express fast and slow rhythmic activities in the stomatogastric ganglion. Journal of Neurophysiology, 95(6):3617–3632.

Buchholtz, F., Golowasch, J., Epstein, I. R., and Marder, E. (1992). Mathematical model of an identified stomatogastric ganglion neuron. Journal of Neurophysiology, 67(2):332–340.

Caplan, J. S., Williams, A. H., and Marder, E. (2014). Many parameter sets in a multicompartment model oscillator are robust to temperature perturbations. Journal of Neuroscience, 34(14):4963–4975.

Connor, J. (1975). Neural repetitive firing: a comparative study of membrane properties of crustacean walking leg axons. Journal of Neurophysiology, 38(4):922–932.

Drion, G., O’Leary, T., and Marder, E. (2015). Ion channel degeneracy enables robust and tunable neuronal firing rates. Proceedings of the National Academy of Sciences, 112(38):E5361–E5370.

Fitzhugh, R. (1966). Theoretical effect of temperature on threshold in the hodgkin-huxley nerve model. The Journal of General Physiology, 49(5):989–1005.

Frankenhaeuser, B. and Moore, L. (1963). The effect of temperature on the sodium and potassium permeability changes in myelinated nerve fibres of xenopus laevis. The Journal of Physiology, 169(2):431–437.

Garrett, S. and Rosenthal, J. J. (2012). RNA editing underlies temperature adaptation in K+ channels from polar octopuses. Science, 335(6070):848–851.

Goaillard, J.-M., Taylor, A. L., Schulz, D. J., and Marder, E. (2009). Functional consequences of animal-to-animal variation in circuit parameters. Nature Neuroscience, 12(11):1424.

Goldman, M. S., Golowasch, J., Marder, E., and Abbott, L. (2001). Global structure, robustness, and modulation of neuronal models. Journal of Neuroscience, 21(14):5229–5238.

Golowasch, J. (2019). Neuronal homeostasis: Voltage brings it all together. Current Biology, 29(13):R641–R644.

Golowasch, J., Abbott, L., and Marder, E. (1999a). Activity-dependent regulation of potassium currents in an identified neuron of the stomatogastric ganglion of the crab cancer borealis. Journal of Neuroscience, 19(20):RC33–RC33.

Golowasch, J., Buchholtz, F., Epstein, I. R., and Marder, E. (1992). Contribution of individual ionic currents to activity of a model stomatogastric ganglion neuron. Journal of Neurophysiology, 67(2):341–349.

Golowasch, J., Casey, M., Abbott, L., and Marder, E. (1999b). Network stability from activity-dependent regulation of neuronal conductances. Neural Computation, 11(5):1079–1096.

Golowasch, J. and Marder, E. (1992). Ionic currents of the lateral pyloric neuron of the stomatogastric ganglion of the crab. Journal of Neurophysiology, 67(2):318–331.

Guttman, R. et al. (1962). Effect of temperature on the potential and current thresholds of axon membrane. The Journal of General Physiology, 46(2):257–266.

Guttman, R. et al. (1966). Temperature characteristics of excitation in space-clamped squid axons. The Journal of General Physiology, 49(5):1007–1018.

Haddad, S. A. and Marder, E. (2018). Circuit robustness to temperature perturbation is altered by neuromodulators. Neuron, 100(3):609–623.

Haley, J. A., Hampton, D., and Marder, E. (2018). Two central pattern generators from the crab, *Cancer borealis*, respond robustly and differentially to extreme extracellular ph. eLife, 7:e41877.

Hamood, A. W., Haddad, S. A., Otopalik, A. G., Rosenbaum, P., and Marder, E. (2015). Quantitative reevaluation of the effects of short-and long-term removal of descending modulatory inputs on the pyloric rhythm of the crab, *Cancer borealis*. eNeuro, 2(1).

Harris-Warrick, R. M., Coniglio, L. M., Barazangi, N., Guckenheimer, J., and Gueron, S. (1995). Dopamine modulation of transient potassium current evokes phase shifts in a central pattern generator network. Journal of Neuroscience, 15(1):342–358.

Heitler, W., Goodman, C. S., and Rowell, C. F. (1977). The effects of temperature on the threshold of identified neurons in the locust. Journal of Comparative Physiology A: Neuroethology, Sensory, Neural, and Behavioral Physiology, 117(2):163–182.

Holland, J. H. (1992). Genetic algorithms. Scientific American, 267(1):66–73.

Hooper, S. L. (1997). Phase maintenance in the pyloric pattern of the lobster *(Panulirus interruptus*) stomatogastric ganglion. Journal of Computational Neuroscience, 4(3):191–205.

Klee, M., Pierau, F.-K., and Faber, D. (1974). Temperature effects on resting potential and spike parameters of cat motoneurons. Experimental Brain Research, 19(5):478–492.

Krupczynski, P. and Schuster, S. (2013). Precision of archerfish c-starts is fully temperature compensated. Journal of Experimental Biology, 216(18):3450–3460.

Kukita, F. (1982). Properties of sodium and potassium channels of the squid giant axon far below 0°c. The Journal of Membrane Biology, 68(1):151–160.

LeMasson, G., Marder, E., and Abbott, L. (1993). Activity-dependent regulation of conductances in model neurons. Science, 259(5103):1915–1917.

Liu, Z., Golowasch, J., Marder, E., and Abbott, L. (1998). A model neuron with activity-dependent conductances regulated by multiple calcium sensors. Journal of Neuroscience, 18(7):2309–2320.

Ma, M. and Koester, J. (1996). The role of K+ currents in frequency-dependent spike broadening in *Aplysia* R20 neurons: a dynamic-clamp analysis. Journal of Neuroscience, 16(13):4089–4101.

Malashchenko, T., Shilnikov, A., and Cymbalyuk, G. (2011). Six types of multistability in a neuronal model based on slow calcium current. PloS One, 6(7):e21782.

Manor, Y., Nadim, F., Abbott, L., and Marder, E. (1997). Temporal dynamics of graded synaptic transmission in the lobster stomatogastric ganglion. Journal of Neuroscience, 17(14):5610–5621.

Marder, E. and Bucher, D. (2007). Understanding circuit dynamics using the stomatogastric nervous system of lobsters and crabs. Annu. Rev. Physiol., 69:291–316.

Marin, B., Barnett, W. H., Doloc-Mihu, A., Calabrese, R. L., and Cymbalyuk, G. S. (2013). High prevalence of multistability of rest states and bursting in a database of a model neuron. PLoS Computational Biology, 9(3):e1002930.

Maynard, D. M. (1972). Simpler networks. Annals of the New York Academy of Sciences, 193(1):59–72.

Nadim, F. and Manor, Y. (2000). The role of short-term synaptic dynamics in motor control. Current Opinion in Neurobiology, 10(6):683–690.

Nadim, F., Manor, Y., Kopell, N., and Marder, E. (1999). Synaptic depression creates a switch that controls the frequency of an oscillatory circuit. Proceedings of the National Academy of Sciences, 96(14):8206–8211.

O’Leary, T. and Marder, E. (2016). Temperature-robust neural function from activity-dependent ion channel regulation. Current Biology, 26(21):2935–2941.

O’Leary, T., Williams, A. H., Caplan, J. S., and Marder, E. (2013). Correlations in ion channel expression emerge from homeostatic tuning rules. Proceedings of the National Academy of Sciences, 110(28):E2645–E2654.

O’Leary, T., Williams, A. H., Franci, A., and Marder, E. (2014). Cell types, network homeostasis, and pathological compensation from a biologically plausible ion channel expression model. Neuron, 82(4):809–821.

Press, W. H., Teukolsky, S. A., Vetterling, W. T., and Flannery, B. P. (1988). Numerical recipes in C. Cambridge University Press, 1:3.

Prinz, A. A., Bucher, D., and Marder, E. (2004). Similar network activity from disparate circuit parameters. Nature Neuroscience, 7(12):1345.

Ratliff, J., Marder, E., and O’Leary, T. (2018). Neural circuit robustness to acute, global physiological perturbations. BioRxiv, page 480830.

Rieder, L. E., Savva, Y. A., Reyna, M. A., Chang, Y.-J., Dorsky, J. S., Rezaei, A., and Reenan, R. A. (2015). Dynamic response of RNA editing to temperature in *Drosophila*. BMC Biology, 13(1):1.

Rinberg, A., Taylor, A. L., and Marder, E. (2013). The effects of temperature on the stability of a neuronal oscillator. PLoS Computational Biology, 9(1):e1002857.

Robertson, R. M. and Money, T. G. (2012). Temperature and neuronal circuit function: compensation, tuning and tolerance. Current Opinion in Neurobiology, 22(4):724–734.

Roemschied, F. A., Eberhard, M. J., Schleimer, J.-H., Ronacher, B., and Schreiber, S. (2014). Cell-intrinsic mechanisms of temperature compensation in a grasshopper sensory receptor neuron. eLife, 3:e02078.

Rosenthal, J. J. (2015). The emerging role of RNA editing in plasticity. Journal of Experimental Biology, 218(12):1812–1821.

Ruff, R. L. (1999). Effects of temperature on slow and fast inactivation of rat skeletal muscle *Na+* channels. American Journal of Physiology-Cell Physiology, 277(5):C937–C947.

Santin, J. M. and Schulz, D. J. (2019). Membrane voltage is a direct feedback signal that influences correlated ion channel expression in neurons. Current Biology, 29(10):1683–1688.

Schauf, C. (1973). Temperature dependence of the ionic current kinetics of *Myxicola* giant axons. The Journal of Physiology, 235(1):197–205.

Schulz, D. J., Goaillard, J.-M., and Marder, E. (2006). Variable channel expression in identified single and electrically coupled neurons in different animals. Nature Neuroscience, 9(3):356.

Schulz, D. J., Goaillard, J.-M., and Marder, E. E. (2007). Quantitative expression profiling of identified neurons reveals cell-specific constraints on highly variable levels of gene expression. Proceedings of the National Academy of Sciences, 104(32):13187–13191.

Sjodin, R. and Mullins, L. (1958). Oscillatory behavior of the squid axon membrane potential. The Journal of General Physiology, 42(1):39–47.

Soofi, W., Goeritz, M. L., Kispersky, T. J., Prinz, A. A., Marder, E., and Stein, W. (2014). Phase maintenance in a rhythmic motor pattern during temperature changes *in vivo*. Journal of Neurophysiology, 111(12):2603–2613.

Soto-Trevino, C., Rabbah, P., Marder, E., and Nadim, F. (2005). Computational model of electrically coupled, intrinsically distinct pacemaker neurons. Journal of Neurophysiology, 94(1):590–604.

Tang, L. S., Goeritz, M. L., Caplan, J. S., Taylor, A. L., Fisek, M., and Marder, E. (2010). Precise temperature compensation of phase in a rhythmic motor pattern. PLoS Biology, 8(8):e1000469.

Tang, L. S., Taylor, A. L., Rinberg, A., and Marder, E. (2012). Robustness of a rhythmic circuit to short-and long-term temperature changes. Journal of Neuroscience, 32(29):10075–10085.

Taylor, B. and Kerkut, G. (1958). The effect of temperature changes on the activity of poikilotherms. Behaviour, 13(3-4):259–279.

Temporal, S., Lett, K. M., and Schulz, D. J. (2014). Activity-dependent feedback regulates correlated ion channel mrna levels in single identified motor neurons. Current Biology, 24(16):1899–1904.

Tran, T., Unal, C. T., Severin, D., Zaborszky, L., Rotstein, H. G., Kirkwood, A., and Golowasch, J. (2019). Ionic current correlations are ubiquitous across phyla. Scientific Reports, 9(1):1687.

Turrigiano, G., Abbott, L., and Marder, E. (1994). Activity-dependent changes in the intrinsic properties of cultured neurons. Science, 264(5161):974–977.

Turrigiano, G., LeMasson, G., and Marder, E. (1995). Selective regulation of current densities underlies spontaneous changes in the activity of cultured neurons. Journal of Neuroscience, 15(5):3640–3652.

